# SARS-CoV-2 immune evasion by variant B.1.427/B.1.429

**DOI:** 10.1101/2021.03.31.437925

**Authors:** Matthew McCallum, Jessica Bassi, Anna De Marco, Alex Chen, Alexandra C. Walls, Julia Di Iulio, M. Alejandra Tortorici, Mary-Jane Navarro, Chiara Silacci-Fregni, Christian Saliba, Maria Agostini, Dora Pinto, Katja Culap, Siro Bianchi, Stefano Jaconi, Elisabetta Cameroni, John E. Bowen, Sasha W Tilles, Matteo Samuele Pizzuto, Sonja Bernasconi Guastalla, Giovanni Bona, Alessandra Franzetti Pellanda, Christian Garzoni, Wesley C. Van Voorhis, Laura E. Rosen, Gyorgy Snell, Amalio Telenti, Herbert W. Virgin, Luca Piccoli, Davide Corti, David Veesler

## Abstract

SARS-CoV-2 entry is mediated by the spike (S) glycoprotein which contains the receptor-binding domain (RBD) and the N-terminal domain (NTD) as the two main targets of neutralizing antibodies (Abs). A novel variant of concern (VOC) named CAL.20C (B.1.427/B.1.429) was originally detected in California and is currently spreading throughout the US and 29 additional countries. It is unclear whether antibody responses to SARS-CoV-2 infection or to the prototypic Wuhan-1 isolate-based vaccines will be impacted by the three B.1.427/B.1.429 S mutations: S13I, W152C and L452R. Here, we assessed neutralizing Ab responses following natural infection or mRNA vaccination using pseudoviruses expressing the wildtype or the B.1.427/B.1.429 S protein. Plasma from vaccinated or convalescent individuals exhibited neutralizing titers, which were reduced 3-6 fold against the B.1.427/B.1.429 variant relative to wildtype pseudoviruses. The RBD L452R mutation reduced or abolished neutralizing activity of 14 out of 35 RBD-specific monoclonal antibodies (mAbs), including three clinical-stage mAbs. Furthermore, we observed a complete loss of B.1.427/B.1.429 neutralization for a panel of mAbs targeting the N-terminal domain due to a large structural rearrangement of the NTD antigenic supersite involving an S13I-mediated shift of the signal peptide cleavage site. These data warrant closer monitoring of signal peptide variants and their involvement in immune evasion and show that Abs directed to the NTD impose a selection pressure driving SARS-CoV-2 viral evolution through conventional and unconventional escape mechanisms.

## Introduction

Coronavirus disease 2019 (COVID-19) is caused by SARS-CoV-2 and is associated with acute respiratory distress syndrome (ARDS), but also with extra-pulmonary complications such as vascular thrombosis, coagulopathy, and a hyperinflammatory syndrome contributing to disease severity and mortality. SARS-CoV-2 infects target cells via the spike glycoprotein (S) that is organized as a homotrimer wherein each monomer is comprised of an S_1_ and an S_2_ subunit(*1*, *2*). The S_1_ subunit comprises the receptor-binding domain (RBD) and the N-terminal domain (NTD) as well as two other domains designated C and D(*3*, *4*). The RBD interacts with the angiotensin-converting enzyme 2 (ACE2) entry receptor on host cells through a subset of RBD amino acids designated the receptor binding motif (RBM)(*1*, *2*, *5*–*7*). The NTD was suggested to bind DC-SIGN, L-SIGN, and AXL which may act as auxiliary receptors(*8*, *9*). Both the RBD and the NTD are targeted by neutralizing antibodies (Abs) in infected or vaccinated individuals and a subset of RBD-specific mAbs is currently being evaluated in clinical trials or are authorized for use in COVID-19 patients (*10*–*22*). The S_2_ subunit is the fusion machinery that merges viral and host membranes to initiate infection and is the target of Abs cross-reacting with multiple coronavirus subgenera due to its higher sequence conservation compared to the S_1_ subunit(*23*–*25*).

The ongoing global spread of SARS-CoV-2 led to the emergence of a large number of viral lineages worldwide, including several variants of concern (VOC). Specifically, the B.1.1.7, B.1.351, and P.1 lineages that originated in the UK, South Africa, and Brazil, respectively, are characterized by the accumulation of a large number of mutations in the spike as well as in other genes(*26*–*28*). Some of these mutations lead to significant reductions in the neutralization potency of NTD- and RBD-specific mAbs, convalescent sera and Pfizer/BioNTech BNT162b2- or Moderna mRNA-1273-elicited sera(*19*, *29*, *30*). The B.1.1.7 variant is on track to become dominant worldwide due to its higher transmissibility(*28*), underscoring the importance of studying and understanding the consequences of SARS-CoV-2 antigenic drift.

The SARS-CoV-2 B.1.427/B.1.429 variant originated in California in May 2020 and has been detected in more than 29 countries to date(*31*, *32*). It is characterized by the S13I, W152C mutations in the NTD and by the L452R mutation in the RBD. The fast rise in the number of cases associated with the B.1.427/B.1.429 lineages led to their classification as a VOC by the US Center for Disease Control (https://www.cdc.gov/coronavirus/2019-ncov/cases-updates/variant-surveillance/variant-info.html).

## Results

### The prevalence of B.1.427/B.1.429 lineages is increasing exponentially

The novel SARS-CoV-2 VOC B.1.427/B.1.429 was reported for the first time at the beginning of 2021 in California(*31*, *33*, *34*). The two lineages B.1.427 and B.1.429 (belonging to clade 20C according to Nextstrain designation) share the same S mutations (S13I, W152C and L452R), but harbor different mutations in other SARS-CoV-2 genes. Molecular clock analysis suggest that the progenitor of both lineages emerged in May 2020, diverging to give rise to the B.1.427 and B.1.429 independent lineages in June-July 2020(*31*). As of March 26, 2021, 4,292 and 10,934 sequenced genomes are reported in GISAID for the B.1.427 and B.1.429 lineages, respectively. These VOCs were detected in California and in other US states, and more recently in 29 additional countries worldwide (**Fig. 1 A to G**). The number of B.1.427/B.1.429 genome sequences deposited increased rapidly since December 2020 (**Fig. 1 B to E**), with a prevalence exceeding 50% in California since February 2021. Collectively, this analysis illustrates the increased prevalence of the B.1.427/B.1.429 VOC, and their progressive geographical spread from California to other US states and countries, which is consistent with the recent finding of their increased transmissibility relative to currently circulating strains(*31*).

**Fig. 1.**
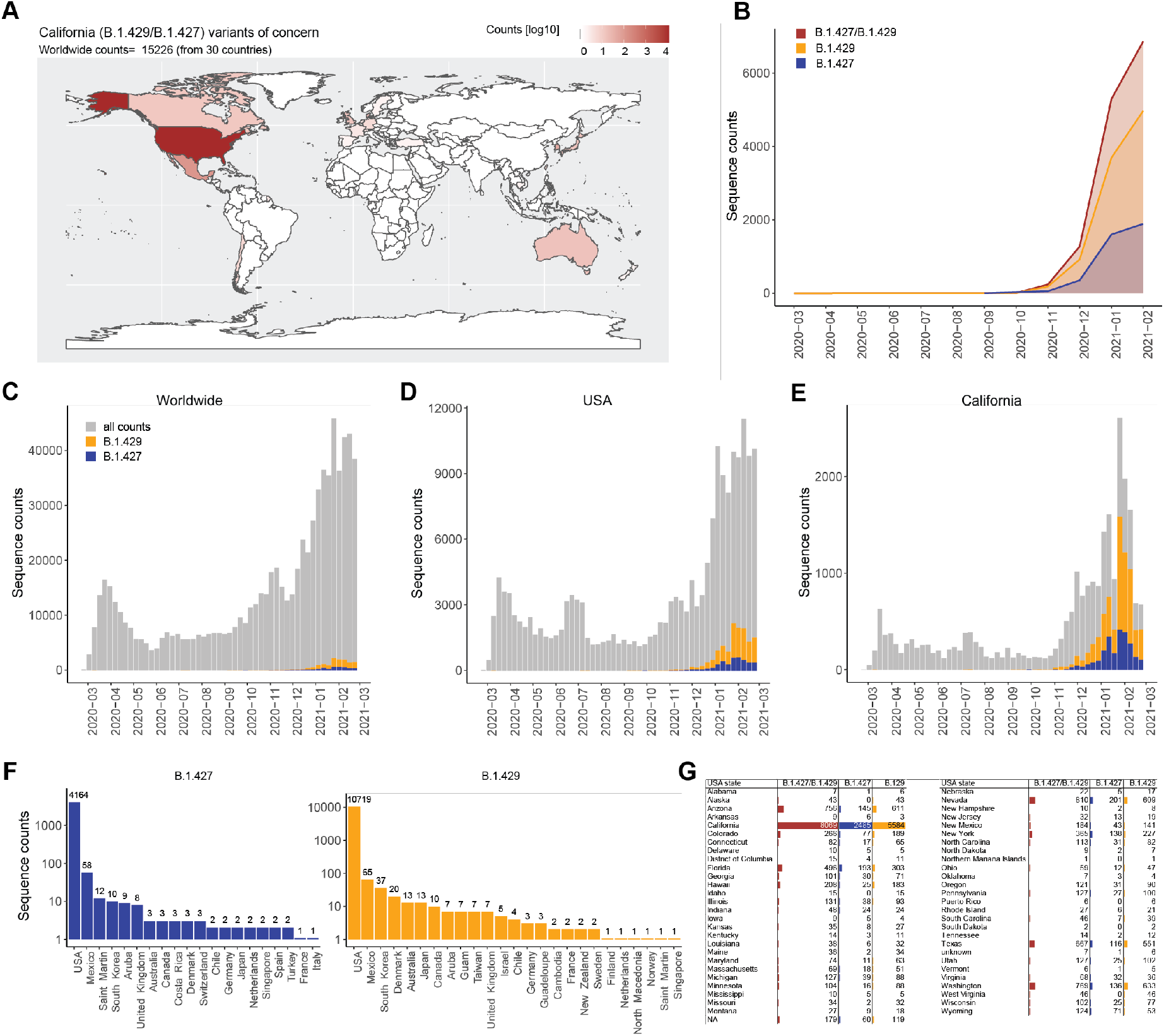
Geographic distribution and evolution of prevalence over time of the SARS-CoV-2 B.1.427/B.1.429 VOC. (**A**) World map showing the geographic distribution and sequence counts of B.1.427/B.1.429 VOC as of March 26, 2021. (**B**) Cumulative and individual B.1.427/B.1.429 VOC sequence counts by month. (**C-E**). Total number of SARS-CoV-2 (grey) and B.1.427/B.1.429 VOC (blue/orange) sequences deposited on a monthly basis worldwide (C), in the US (D) and in California (E). (**F** and **G**) Total number of B.1.427 and B.1.429 sequences deposited by country (F) and by US states (G) as of March 26, 2021.

### B.1.427/ B.1.429 S reduces sensitivity to vaccinees’ plasma

To assess the impact of the three mutations present in the B.1.427/B.1.429 S glycoprotein on neutralization, we first compared side-by-side the neutralization potency of mRNA vaccine-elicited Abs against wildtype (D614G) S and B.1.427/B.1.429 S pseudoviruses. We used plasma from eleven individuals who received two doses of Moderna mRNA-1273 vaccine and from fourteen individuals who received two doses of Pfizer/BioNtech BNT162b2 vaccine collected between 7 and 27 days after booster immunization. All vaccinees had substantial plasma neutralizing activity against wildtype SARS-CoV-2 S pseudotyped viruses. Using a lentiviral (HIV) pseudotyping system, geometric mean titers (GMTs) showed that the average neutralization potency of the Moderna mRNA1273-elicited plasma was reduced 2.8-fold for B.1.427/B.1.429 S (GMT: 204) compared to wildtype (D614G) S (GMT: 573) whereas it was reduced 4-fold with Pfizer/BioNtech BNT162b2-elicited plasma (wildtype GMT: 128 versus B.1.427/B.1.429 GMT: 535) **(Fig. 2A-B)**. Using a vesicular stomatitis virus (VSV) pseudotyping system, we observed a 3-fold average reduction of Pfizer/BioNtech BNT162b2-elicited plasma neutralizing activity against B.1.427/B.1.429 S (GMT: 95) compared to wildtype (D614G) S (GMT: 257) pseudoviruses **(Fig. 2C-D)**. In a parallel analysis, we analyzed 18 individuals, 5 of which were previously infected with SARS-CoV-2, who received two doses of Pfizer/BioNtech BNT162b2 vaccine and whose samples were collected between 14 and 28 days after booster immunization. We compared side-by-side the neutralization potency of Pfizer/BioNtech BNT162b2 vaccine-elicited Abs against wildtype (D614) S, B.1.427/B.1.429 S, as well as B.1.1.7 S, B.1.351 S and P.1 S VSV pseudotyped viruses using Vero E6 expressing TMPRSS2 as target cells. GMTs plasma neutralization potency was reduced 2.8-fold for B.1.427/B.1.429 S (GMT: 248) compared to wildtype (D614) S (GMT: 681), which is a comparable decrease to that observed with B.1.351 (GMT: 211, 3.2-fold reduction) and greater to that observed with B.1.1.7 and P.1 (GMT: 545 and 389, 1.2-fold and 1.7-fold reduction, respectively) pseudotyped viruses **(Fig. 3E-H)**. These data indicate that the B.1.427/B.1.429 S mutations lead to a modest but significant reduction of neutralization potency from vaccine-elicited plasma due to the substitution of one RBD and two NTD residues.

**Fig. 2.**
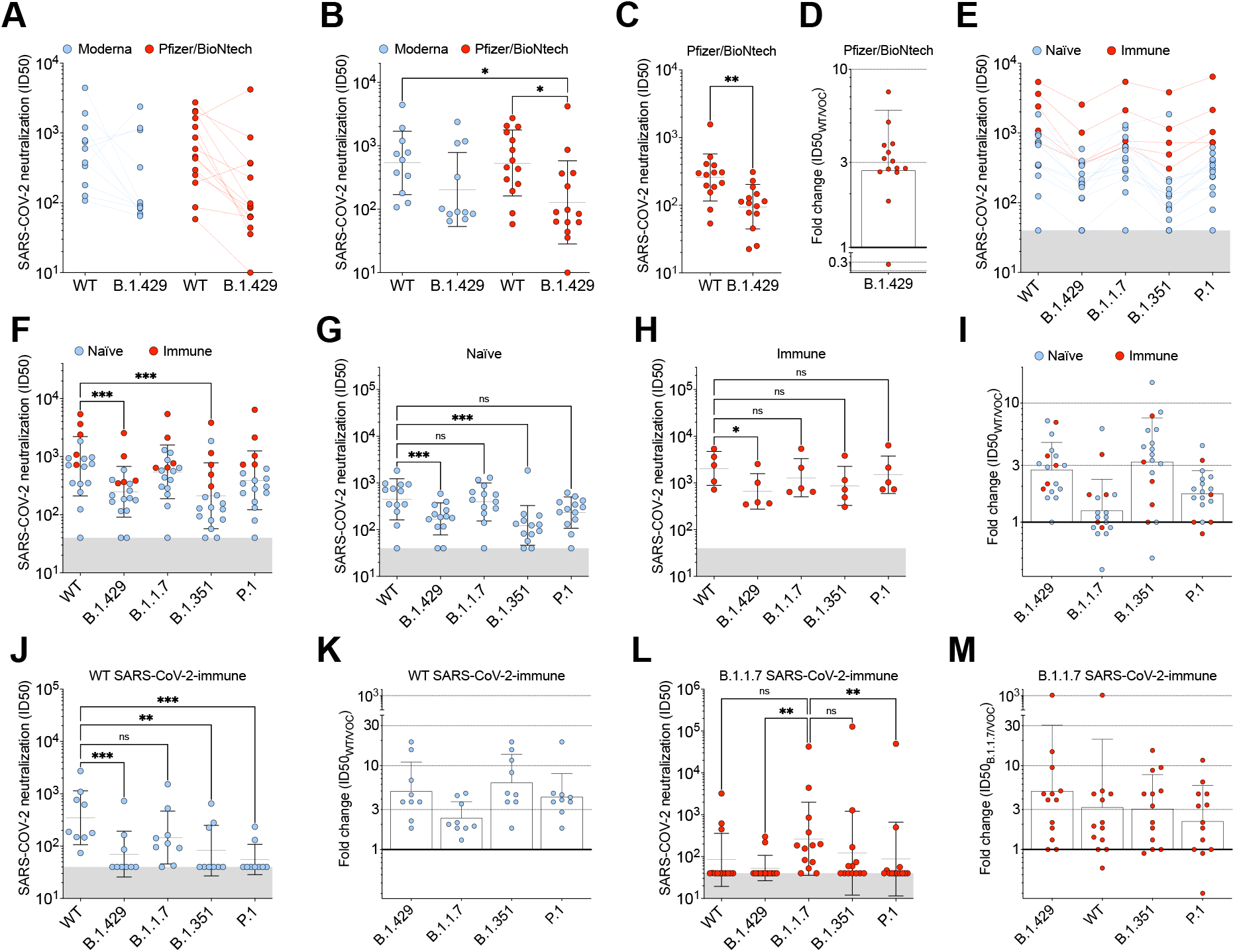
B.1.427/B.1.429 S pseudotyped virus neutralization by vaccine-elicited and COVID-19 convalescent plasma. (**A-B**) Neutralizing Ab titers shown as pairwise connected (**A**) or the geometric mean titer, GMT (**B**) against HIV pseudotyped viruses harboring wildtype (WT) D614G SARS-CoV-2 S or B.1.427/B.1.429 (B.1.429) S determined using plasma from individuals who received two doses of Pfizer/BioNtech BNT162b2 mRNA vaccine (red) or of Moderna mRNA-1273 vaccine (blue). (**C**) Neutralizing Ab GMT against VSV pseudotyped viruses harboring WT D614G SARS-CoV-2 S or B.1.427/B.1.429 S determined using plasma from individuals who received two doses of Pfizer/BioNtech BNT162b2 mRNA vaccine. (**D**) Fold change of neutralizing GMTs compared to wildtype based on (C). (**E-H**) Neutralizing Ab titers (ID50) against VSV pseudotyped viruses harboring WT (D614) SARS-CoV-2 S, B.1.429 S, B.1.1.7 S, B.1.351 S, or P.1 S determined using plasma from naïve (blue, G) and immune (red, H) individuals who received two doses of Pfizer/BioNtech BNT162b2 mRNA vaccine. (**I**) Fold change of neutralizing GMTs compared to wildtype based on (E). (**J-M**) Neutralizing Ab titers against VSV pseudotyped viruses harboring WT (D614) SARS-CoV-2 S, B.1.429 S, B.1.1.7 S, B.1.351 S or P.1 S determined using plasma from convalescent individuals who were infected with wildtype (J, K) or B.1.1.7 SARS-CoV-2 (L, M). Fold change of neutralizing GMTs was compared to wildtype (K) or B.1.1.7 (M). Neutralization data shown in (A-D) and (E-M) performed using 293T-ACE2 and VeroE6-TMPRSS2, respectively.

**Fig. 3.**
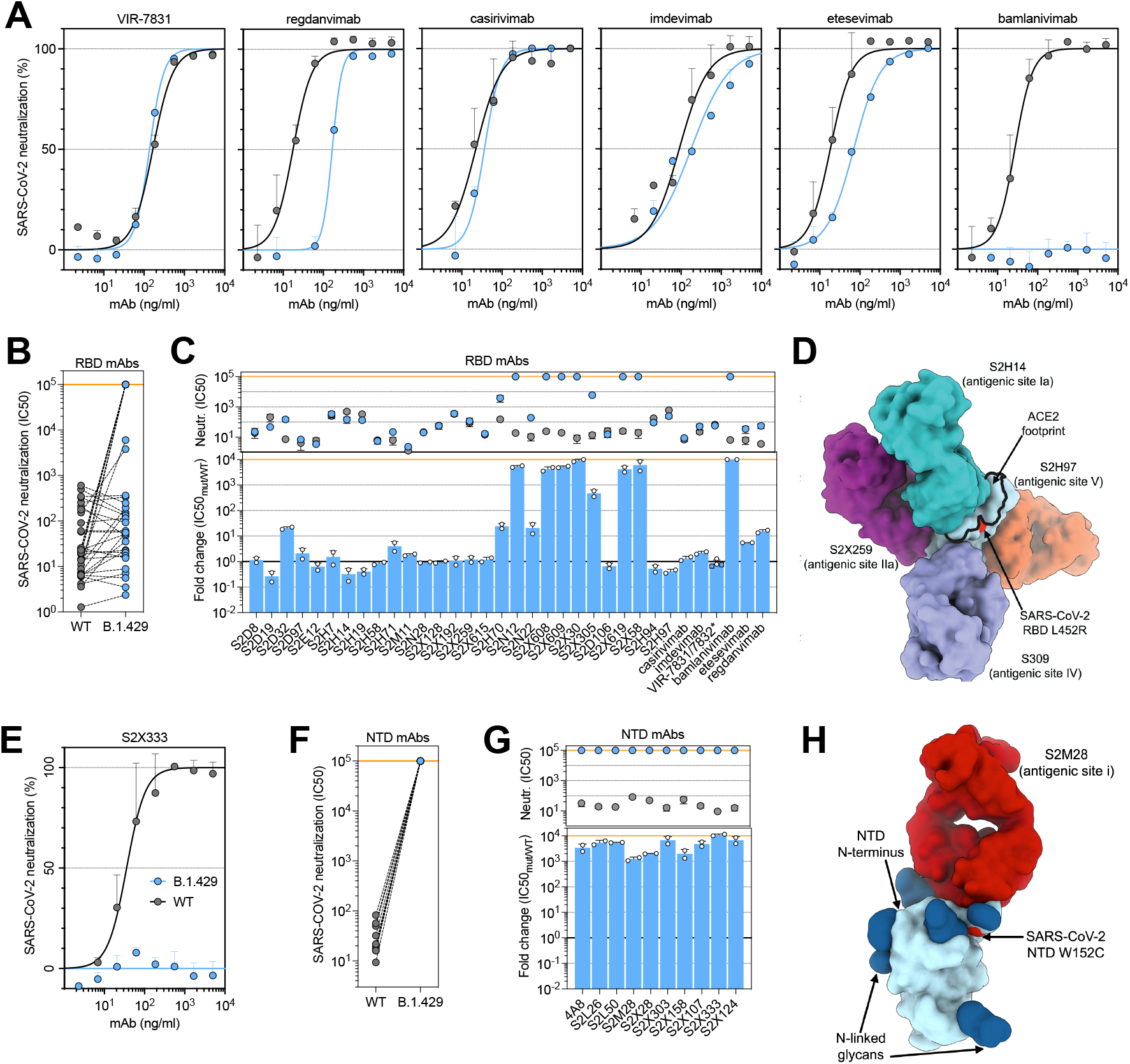
Neutralization by a panel of RBD- and NTD-specific mAbs against wildtype and B.1.427/B.1.429 SARS-CoV-2 S pseudoviruses. (**A,E**) Neutralization of SARS-CoV-2 pseudotyped VSV carrying wild-type (grey) or B.1.427/B.1.429 (blue) S protein by clinical-stage RBD mAbs (A) and an NTD-targeting mAb (S2X333) (E). Data are representative of *n = 2* replicates. (**B,F**) Neutralization of SARS-CoV-2-VSV pseudotypes carrying wildtype or B.1.427/B.1.429 S by 35 mAbs targeting the RBD and 10 mAbs targeting the NTD. Data are the mean of 50% inhibitory concentration (IC_50_) values (ng/ml) of *n = 2* independent experiments. Non-neutralizing IC_50_ titers were set at 10^5^ ng/ml. (**C,G**) Neutralization by RBD-specific (C) and NTD-specific (G) mAbs showed as mean IC_50_ values (top) and mean fold change for B.1.427/B.1.429 S (blue) relative to wildtype (D614G) S (grey) VSV pseudoviruses. VIR-7831 is a derivative of S309 mAb. *, VIR-7832 (variant of VIR-7831 carrying the LS-GAALIE mutations in Fc) shown as squares. Non-neutralizing IC_50_ titers and fold change were set at 10^5^ ng/ml and 10^4^, respectively. (**D**) Surface representation of the SARS-CoV-2 RBD bound to the S2H14 (antigenic site Ia), S2X259 (antigenic site IIa), S309 (antigenic site IV) and S2H97 (antigenic site V) Fab fragments shown as surfaces. (**H**) Surface representation of the SARS-CoV-2 NTD bound to the S2M28 (antigenic site i) Fab fragment shown as surface.

We also analyzed plasma from 9 convalescent donors, who experienced symptomatic COVID-19 in early 2020 (likely exposed to the Wuhan-1 or a closely related SARS-CoV-2 isolate) collected 15 to 28 days after symptom onset, and 13 additional plasma samples from healthcare workers collected approximately 1 month after infection with the B.1.1.7 VOC during an outbreak occurring in a nursing home in January 2021 in Ticino, Switzerland. The neutralization potency of the 9 convalescent donor plasma was reduced 4.9-fold for B.1.427/B.1.429 S (GMT: 70) compared to wildtype (D614G) S (GMT: 348), similar to what we observed with B.1.351 (6.2-fold, GMT: 82) and P.1 (4.2-fold, GMT: 55) pseudotyped viruses **(Fig. 2I-K)**. In several cases the level of neutralizing activity against the VOC was found to be below the limit of detection. The plasma neutralizing activity of the 13 healthcare workers infected with B.1.1.7 VOC was found to be highest against the homotypic pseudovirus (i.e., B.1.1.7, GMT: 265) and to be reduced to undetectable levels in most cases against the wildtype (D614G) pseudovirus and the other VOC tested, including B.1.427/B.1.429 **(Fig. 2 I,L-M)**, which is different from what was observed with B.1.351-elicited Abs (*35*, *36*). Since B.1.1.7 is now the prevalent lineage in Europe and several other countries(*26*) the finding that infection with B.1.1.7 elicits a low level of cross-neutralizing Abs towards the other co-circulating lineages is concerning.

These findings show that the three mutations present in the B1.427/B.1.429 S glycoprotein decrease the neutralizing activity of vaccine-elicited and infection-elicited Abs, suggesting that these lineage-defining residue substitutions are associated with immune evasion. However, the data also underscore the higher quality of Ab responses induced by vaccination compared to infection and their enhanced resilience to mutations against all VOC.

### B.1.427/B.1.429 S mutations reduce sensitivity to RBD- and NTD-specific antibodies

To evaluate the contribution of RBD and NTD substitutions to the reduced neutralization potency of vaccinees and convalescent plasma, we compared the neutralizing activity of 34 RBD and 10 NTD mAbs against the wildtype (D614 S or B.1.427/B.1.429 S variant using a VSV pseudotyping system(*1*, *37*).

We used a panel of RBD-specific mAbs (including 6 clinical mAbs) recognizing distinct antigenic sites spanning the receptor-binding motif (RBM, antigenic sites Ia and Ib), the cryptic antigenic site II, the exposed, N343 glycan-containing antigenic site IV and the newly discovered, cryptic antigenic site V(*10*, *11*). A total of 14 out of 35 mAbs showed a reduced neutralization potency when comparing B.1.427/B.1.429 S and wildtype (D614G) S pseudoviruses **(Fig. 3A-D** and **Supplemental Fig. 2)**. Regdanvimab (CT-P59), and to a smaller extent etesevimab, showed a reduction in neutralization potency, whereas bamlanivimab (LY-CoV555) entirely lost its neutralizing activity due to the central location of L452R in the epitopes recognized by these mAbs **(Supplemental Fig. 1)**. Neutralization mediated by the casirivimab/imdevimab mAb cocktail (REGN10933 and REGN10987)(*14*, *15*), which received an emergency use authorization in the US, and by VIR-7831 mAb(*10*, *38*) (derivative of S309), which recently was shown to provide 85% protection against hospitalization and deaths in the COMET clinical trial, is unaffected by the L452R mutation. To address the role of B.1.427/B.1.429 L452R mutation in the neutralization escape from RBD-specific antibodies, we tested the binding of 35 RBD-specific mAbs to WT and L452R mutant RBD by biolayer interferometry (**Supplemental Fig. 3)**. The 10 RBD-specific mAbs that showed at least 10-fold reduced neutralization of B.1.427/B.1.429 variant were also found to poorly bind to L452R RBD mutant, demonstrating a role for this mutation as an escape mechanism for certain RBD-targeting mAbs. Moreover, we found that the neutralizing activity of all NTD-specific neutralizing mAbs tested was abolished as a result of the presence of the S13I and W152C mutations **(Fig. 3E-H)**. These data suggest that the decreased potency of neutralization of the B.1.427/B.1.429 variant observed with vaccine-elicited and infection-elicited plasma results from evasion of both RBD- and NTD-specific mAb-mediated neutralization.

### S13I-mediated immune evasion of B.1.427/B.1.429

To reveal the molecular basis of the observed loss of NTD-directed mAb neutralizing activity, we analyzed binding of a panel of NTD-specific mAbs to recombinant SARS-CoV-2 NTD variants using ELISA. The S13I mutation dampened binding of 5 mAbs and abrogated binding of 5 additional mAbs out of 11 neutralizing mAbs evaluated **(Figure. 4A** and **Supplemental Fig. 4).** Furthermore, the W152C mutation reduced recognition of six NTD neutralizing mAbs, including a complete loss of binding for two of them, with a complementary pattern to that observed for S13I **(Fig. 4A** and **Supplemental Fig. 4)**. The B.1.427/B.1.429 S13I/W152C NTD did not bind to any NTD-directed neutralizing mAbs, which are known to target a single antigenic site (antigenic site i)(*12*), whereas binding of the non-neutralizing S2L20 mAb to the NTD antigenic site iv was not affected by any mutants, confirming proper retention of folding **(Fig. 4A and Supplemental Fig. 4)**.

**Fig. 4.**
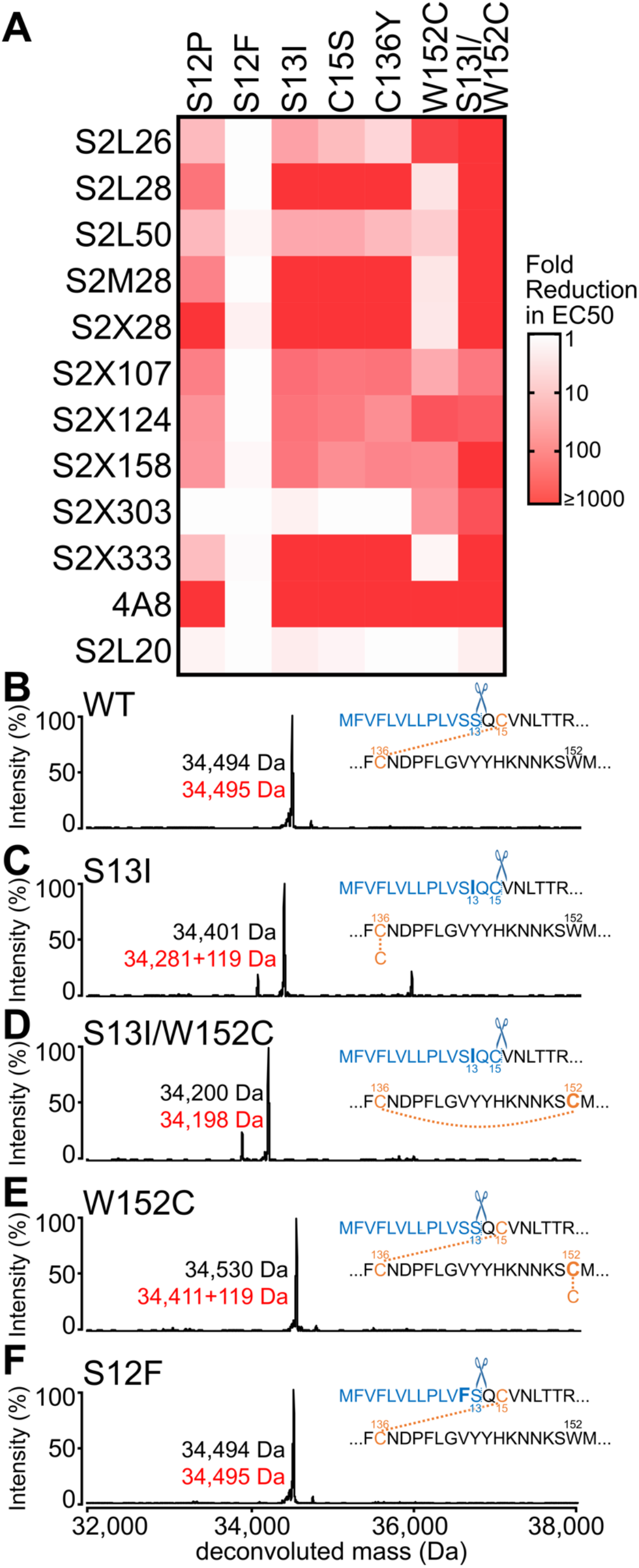
The B.1.427/B.1.429 S S13I signal peptide mutation leads to immune evasion. (**A**) Binding of a panel of 11 neutralizing (antigenic site i) and 1 non-neutralizing (antigenic site iv) NTD-specific mAbs to recombinant SARS-CoV-2 NTD variants analyzed by ELISA displayed as a heat map. (**B-F**) Deconvoluted mass spectra of purified NTD constructs, including the wildtype NTD with the native signal peptide (B), the S13I NTD (C), the S13I and W152C NTD (D), the W152C NTD (E), and the S12F NTD (F). The empirical mass (black) and theoretical mass (red) are shown beside the corresponding peak. Additional 119 Da were observed for the S13I and W152C NTDs corresponding to cysteinylation (119 Da) of the free cysteine residue in these constructs (as L-cysteine was present in the expression media). The cleaved signal peptide (blue text) and subsequent residue sequence (black text) are also shown based on the MS results. Mutated residues are shown in bold. Cysteines are highlighted in light orange (unless in the cleaved signal peptide) while disulfide bonds are shown as dotted light orange lines between cysteines. Residues are numbered for reference.

We previously showed that disruption of the C15/C136 disulfide bond that connects the N-terminus to the rest of the NTD, through mutation of either residue or alteration of the signal peptide cleavage site, abrogates the neutralizing activity of mAbs targeting the NTD antigenic supersite (site i)(*12*). As the S13I substitution resides in the signal peptide and is predicted to shift the signal peptide cleavage site from S13-Q14 to C15-V16, we hypothesized that this substitution indirectly affects the integrity of NTD antigenic site i, which comprises the N-terminus. Mass spectrometry analysis of the S13I and S13I/W152C NTD variants confirmed that signal peptide cleavage occurs immediately after residue C15 **(Figure. 4B-D).** As a result, C136, which would otherwise be disulfide linked to C15, is cysteinylated in the S13I NTD **(Fig. 4C and Supplemental Fig. 5)**. Likewise, the W152C mutation, which also introduces a free cysteine, was also found to be cysteinylated in the W152C NTD **(Fig. 4E).** Notably, dampening of NTD-specific neutralizing mAb binding is even stronger for the S13I mutant than for the S12P mutant **(Fig. 4A)**. Conversely, we did not observe any effect on mAb binding of the S12F substitution, which has also been detected in clinical isolates, in agreement with the fact that it did not affect the native signal peptide cleavage site (i.e. it occurs at the native S13-Q14 position), as observed by mass spectrometry **(Fig. 4F)**. While the S13I and W152C NTD variants were respectively cysteinylated at positions C136 and W152C, due to the presence of unpaired cysteines, the double mutant S13I/W152C was not cysteinylated, suggesting that C136 and W152C had formed a disulfide bond with each other **(Fig. 4C-E).** Tandem mass-spectrometry analysis of non-reduced, digested peptides identified linked discontinuous peptides containing C136 and W152C **(Supplemental Fig. 5)**. The predominant observation of these two residues in the set of digested peptides thereby confirmed that a disulfide bond forms between C136 and W152C in the S13I/W152C NTD of the B.1.427/B.1.429 variant.

Collectively, these findings demonstrate that the S13I and W152C mutations found in the B.1.427/B.1.429 S variant are jointly responsible for escape from NTD-specific mAbs, due to deletion of the SARS-CoV-2 S two N-terminal residues and overall rearrangement of the NTD antigenic supersite. Our data support that the SARS-CoV-2 NTD evolved a previously undescribed compensatory mechanism to form an alternative disulfide bond and that mutations of the S signal peptide occur *in vivo* in a clinical setting to promote immune evasion.

## Discussion

Serum or plasma neutralizing activity is a correlate of protection against SARS-CoV-2 challenge in non-human primates(*39*, *40*) and several monoclonal neutralizing Abs have demonstrated their ability to reduce viral burden as well as to decrease hospitalization and mortality in clinical trials(*10*, *14*, *15*, *22*, *38*, *41*). The data presented here indicate that SARS-CoV-2 B.1.427/B.1.429 is associated with a reduction of sensitivity to plasma neutralizing Abs elicited by vaccination with two doses of Pfizer/BioNTech BNT162b2 or Moderna mRNA-1273 and by infection with the prototypic SARS-CoV-2 and the B.1.1.7 VOC. This reduction correlates with the loss of neutralizing activity observed with all human mAbs directed to the NTD evaluated as well as reduction or loss of inhibition for about a third of RBD-specific mAbs tested. These findings are reminiscent of recent observations made with other VOC, such as the B.1.1.7, B.1.351 and P.1, which also accumulated RBD and NTD mutations negatively affecting the neutralization potency of polyclonal Abs or mAb(*12*, *19*, *29*, *30*, *42*–*44*).

The single L452R mutation present in the SARS-CoV-2 B.1.427/B.1.429 S RBD leads to a reduction or abrogation of the neutralizing activity of 10 out of 34 RBD-specific mAbs evaluated, including regdanvimab (CT-P59), etesevimab (LY-CoV016) and bamlanivimab (LY-CoV555). Conversely, neutralization mediated by the Regeneron casirivimab/imdevimab mAb cocktail and by the VIR-7831/VIR-7832 (derivatives of S309) mAbs was indistinguishable against the wildtype and the B.1.427/B.1.49 variant. The observed L452R-mediated immune evasion of B.1.427/B.1.429 S concurs with previous findings that this substitution reduced the binding or neutralizing activity of some mAbs prior to the description of the B.1.427/B.1.429 variant(*45*–*48*). The acquisition of the L452R substitution by multiple lineages across multiple continents is suggestive of positive selection, which might result from the selective pressure of RBD-specific neutralizing Abs(*34*). We anticipate that these data will guide public health policies regarding the deployment of these clinical mAbs for use in early therapy of COVID-19.

Whereas RBD neutralizing mAbs target various antigenic sites, all known NTD-specific neutralizing mAbs recognize the same antigenic supersite(*12*, *16*–*18*). Both types of mAbs represent a key aspect of immunity to SARS-CoV-2 influencing viral evolution(*12*). The SARS-CoV-2 NTD undergoes fast antigenic drift and accumulates a larger number of prevalent mutations and deletions relative to other regions of the S glycoprotein(*12*, *49*). For instance, the L18F substitution and the deletion of residue Y144 are found in 8% and 26% of viral genomes sequenced and are present in the B.1.351/P.1 lineages and the B.1.1.7 lineage, respectively. Both of these mutations are associated with reduction or abrogation of mAb binding and neutralization(*12*, *29*). The finding that multiple circulating SARS-CoV-2 variants map to the NTD, including several of them in the antigenic supersite (site i), suggests that the NTD is subject to a strong selective pressure from the host humoral immune response, as supported by the identification of deletions within the NTD antigenic supersite in immunocompromised hosts with prolonged infections(*50*–*52*) and the *in vitro* selection of SARS-CoV-2 S escape variants with NTD mutations that decrease binding and neutralization potency of COVID-19 convalescent patient sera or mAbs(*12*, *29*, *53*, *54*).

The SARS-CoV-2 B.1.427/B.1.429 S NTD harbors the S13I and W152C substitutions and we demonstrate here that the former mutation leads to shifting the signal peptide cleavage site, effectively deleting the first two amino acid residues of the S glycoprotein (Q14 and C15). This deletion disrupts the C15/C136 disulfide bond that staples the N-terminus to the rest of the NTD galectin-like β-sandwich and thereby compromises the integrity of the NTD site of vulnerability. The SARS-CoV-2 B.1.427/B.1.429 S variant therefore relies on an indirect and unusual neutralization-escape strategy.

The S13I/W152C mutations are efficiently evading the neutralizing activity of NTD-specific mAbs, and the acquisition of additional RBD mAb escape mutations (in addition to L452R) could further dampen Ab-mediated SARS-CoV-2 neutralization for B.1.427/B.1.429. For example, the independent acquisition of the E484K mutation in the B.1.351, P.1, B.1.526 variants and more recently the B.1.1.7 variant(*29*) suggests this could also occur in the B.1.427/B.1.429 lineages, as supported by the presence in GISAID of 4 genome sequences with the E484K RBD mutation in the B.1.427 variant. Alternatively, the S13I mutation could emerge in any of these variants. We note that the S13I mutation was also detected in the SARS-CoV-2 B.1.526 lineage, which was originally described in New York(*55*, *56*). Understanding the newfound mechanism of immune evasion of the emerging variants, such as the signal peptide modification described herein, is as important as sequence surveillance itself to successfully counter the ongoing pandemic.

## ACKNOWLEDGEMENTS

We thank Hideki Tani (University of Toyama) for providing the reagents necessary for preparing VSV pseudotyped viruses. This study was supported by the National Institute of Allergy and Infectious Diseases (DP1AI158186 and HHSN272201700059C to D.V., and U01 AI151698-01 to WCVV), a Pew Biomedical Scholars Award (D.V.), Investigators in the Pathogenesis of Infectious Disease Awards from the Burroughs Wellcome Fund (D.V.), Fast Grants (D.V.), the Natural Sciences and Engineering Research Council of Canada (M.M.), the Pasteur Institute (M.A.T).

## AUTHOR CONTRIBUTIONS

Conceived study: L.P., D.C., D.V. Designed study and experiments: M.M., J.B., A.D.M, A.C., A.C.W., J.d.I., M.A.T. Performed mutagenesis for mutant expression plasmids: M.M., E.C. and K.C. Performed mutant expression: M.M., J.E.B., E.C. and S.J. Contributed to donor’s recruitment and plasma samples collection: S.B.G., G.B., A.F.P, C.G., S.T., W.V. Produced pseudoviruses and carried out pseudovirus neutralization assays. A.C.W., M.A.T., M.J.N., J.B., A.D.M., D.P., C.S., C.S-F. Bioinformatic analysis: J.d.I and A.T. Analyzed the data and prepared the manuscript with input from all authors: M.M., J.B., A.D.M., L.E.R., G.S., L.P., D.C. and D.V; supervision: M.S.P., L.P., G.S., H.W.V., D.C., and D.V

## MATERIALS AND METHODS

### Cell lines

Cell lines used in this study were obtained from ATCC (HEK293T and Vero E6) or ThermoFisher Scientific (Expi CHO cells, FreeStyle™ 293-F cells and Expi293F™ cells).

### B.1.427/B.1.429 prevalence analysis

The viral sequences and the corresponding metadata were obtained from GISAID EpiCoV project (https://www.gisaid.org/). Analysis was performed on sequences submitted to GISAID up to March 26^th^, 2021. S protein sequences were either obtained directly from the protein dump provided by GISAID or, for the latest submitted sequences that were not incorporated yet in the protein dump at the day of data retrieval, from the genomic sequences with the exonerate(*57*) 2 2.4.0--haf93ef1_3 (https://quay.io/repository/biocontainers/exonerate?tab=tags) using protein to DNA alignment with parameters *-m protein2dna --refine full --minintron 999999 --percent 20* and using accession YP_009724390.1 as a reference. Multiple sequence alignment of all human spike proteins was performed with mafft(*58*) 7.475--h516909a_0 (https://quay.io/repository/biocontainers/mafft?tab=tags) with parameters *--auto --reorder --keeplength --addfragments* using the same reference as above. S sequences that contained >10% ambiguous amino acid or that were < than 80% of the canonical protein length were discarded. A total of 849,975 sequences were used for analysis. Figures were generated with R 4.0.2 () using ggplot 2 3.3.2 and sf 0.9-7 packages

### Sample donors

Samples were obtained from SARS-CoV-2 recovered and vaccinated individuals under study protocols approved by the local Institutional Review Boards (Canton Ticino Ethics Committee, Switzerland,). All donors provided written informed consent for the use of blood and blood components (such as PBMCs, sera or plasma). Samples were collected 14 and 28 days after symptoms onset or after vaccination.

### Ab discovery and recombinant expression

Human mAbs were isolated from plasma cells or memory B cells of SARS-CoV or SARS-CoV-2 immune donors, as previously described(*10*, *59*). Other clinical-stage mAbs (casirivimab, imdevimab, bamlanivimab, etesevimab and regdanbimab) were produced recombinantly based on gene synthesis of VH and VL sequences retrieved from publicly available sequences. Recombinant antibodies were expressed in ExpiCHO cells at 37 °C and 8% CO2. Cells were transfected using ExpiFectamine. Transfected cells were supplemented 1 day after transfection with ExpiCHO Feed and ExpiFectamine CHO Enhancer. Cell culture supernatant was col- lected eight days after transfection and filtered through a 0.2 μm filter. Recombinant antibodies were affinity purified on an ÄKTA xpress FPLC device using 5 mL HiTrapTM MabSelectTM PrismA columns followed by buffer exchange to Histidine buffer (20 mM Histidine, 8% sucrose, pH 6) using HiPrep 26/10 desalting columns.

### Antibody binding measurements using bio-layer interferometry (BLI)

MAbs were diluted to 3 μg/ml in kinetic buffer (PBS supplemented with 0.01% BSA) and immobilized on Protein A Biosensors (FortéBio). Antibody-coated biosensors were incubated for 2 min with a solution containing 5 μg /ml of WT, L452R SARS-CoV-2 RBD in kinetic buffer, followed by a 2-min dissociation step. Change in molecules bound to the biosensors caused a shift in the interference pattern that was recorded in real time using an Octet RED96 system (FortéBio). The binding response over time was used to plot binding data using GraphPad PRISM software (version 9.0.0).

### Serum/plasma and mAbs pseudovirus neutralization assays

#### VSV pseudovirus generation

Replication defective VSV pseudovirus(*60*) expressing SARS-CoV-2 spike proteins corresponding to the different VOC were generated as previously described(*61*) with some modifications. Lenti-X 293T cells (Takara, 632180) were seeded in 10-cm^2^ dishes at a density of 5e6 cells per dish c and the following day transfected with 10 μg of spike expression plasmid with TransIT-Lenti (Mirus, 6600) according to the manufacturer’s instructions. One day post-transfection, cells were infected with VSV-luc (VSV-G) with an MOI of 3 for 1 h, rinsed three times with PBS containing Ca2+/Mg2+, then incubated for an additional 24 h in complete media at 37°C. The cell supernatant was clarified by centrifugation, filtered (0.45 um), aliquoted, and frozen at 80°C.

#### VSV pseudovirus neutralization for the testing of mAbs

Vero-E6 and Vero E6-TMPRSS2 were grown in DMEM supplemented with 10% FBS and seeded into clear bottom white 96 well plates (PerkinElmer, 6005688) at a density of 2e4 cells per well. The next day, mAbs or plasma were serially diluted in pre-warmed complete media, mixed with pseudoviruses and incubated for 1 h at 37°C in round bottom polypropylene plates. Media from cells was aspirated and 50 μl of virus-mAb/plasma complexes were added to cells and then incubated for 1 h at 37°C. An additional 100 μL of prewarmed complete media was then added on top of complexes and cells incubated for an additional 16-24 h. Conditions were tested in triplicate wells on each plate and at least six wells per plate contained untreated infected cells (defining the 0% of neutralization, “MAX RLU” value) and infected cells in the presence of S2E12 and S2X259 at 25 μg/ml each (defining the 100% of neutralization, “MIN RLU” value). Virus-mAb/plasma-containing media was then aspirated from cells and 100 μL of a 1:2 dilution of SteadyLite Plus (Perkin Elmer, 6066759) in PBS with Ca^++^ and Mg^++^ was added to cells. Plates were incubated for 15 min at room temperature and then were analyzed on the Synergy-H1 (Biotek). Average of Relative light units (RLUs) of untreated infected wells (MAX RLU_ave_) was subtracted by the average of MIN RLU (MIN RLU_ave_) and used to normalize percentage of neutralization of individual RLU values of experimental data according to the following formula: (1-(RLU_x_ - MIN RLU_ave_) / (MAX RLU_ave_ – MIN RLU_ave_)) x 100. Data were analyzed and visualized with Prism (Version 9.1.0). IC50 (mAbs) and ID50 (plasma) values were calculated from the interpolated value from the log(inhibitor) versus response – variable slope (four parameters) nonlinear regression with an upper constraint of ≤100, and a lower constrain equal to 0. Each neutralization experiment was conducted on two independent experiments, i.e., biological replicates, where each biological replicate contains a technical triplicate. IC50 values across biological replicates are presented as arithmetic mean ± standard deviation. The loss or gain of neutralization potency across spike variants was calculated by dividing the variant IC50/ID50 by the parental IC50/ID50 within each biological replicate, and then visualized as arithmetic mean ± standard deviation.

#### HIV pseudovirus generation

HIV D614G SARS-CoV-2 S and B.1.427/B.1.429 S pseudotypes were prepared as previously described (*62*, *63*). Briefly, HEK293T cells were co-transfected using Lipofectamine 2000 (Life Technologies) with an S-encoding plasmid, an HIV Gag-Pol, HIV Tat, HIV Rev1B packaging construct, and the HIV transfer vector encoding a luciferase reporter according to the manufacturer’s instructions. Cells were washed 3 × with Opti-MEM and incubated for 5 h at 37°C with transfection medium. DMEM containing 10% FBS was added for 60 h. The supernatants were harvested by spinning at 2,500 g, filtered through a 0.45 μm filter, concentrated with a 100 kDa membrane for 10 min at 2,500 g and then aliquoted and stored at −80°C.

HEK203-hACE2 cells were cultured in DMEM with 10% FBS (Hyclone) and 1% PenStrep with 8% CO_2_ in a 37°C incubator (ThermoFisher). One day or more prior to infection, 40 μL of poly-lysine (Sigma) was placed into 96-well plates and incubated with rotation for 5 min. Poly-lysine was removed, plates were dried for 5 min then washed 1 × with water prior to plating cells. The following day, cells were checked to be at 80% confluence. In a half-area 96-well plate a 1:3 serial dilution of HCP was made in DMEM in 22 μL final volume. 22 μL of diluted pseudovirus was then added to the serial dilution and incubated at room temperature for 30-60 min. Mixture was then added to cells for two hours. Following the two hour incubation, 44μL of DMEM with 20%FBS (Hyclone) and 2% PenStrep was added and incubated for 48 hours. After 48 hours, 40 μL/well of One-Glo-EX substrate (Promega) was added to the cells and incubated in the dark for 5-10 min prior reading on a BioTek plate reader. Measurements were done in at least duplicate. Relative luciferase units were plotted and normalized in Prism (GraphPad) using as zero value cells alone or infected with virus alone as 100%. Nonlinear regression of log(inhibitor) versus normalized response was used to determine IC_50_ values from curve fits. Kruskal Wallis tests were used to compare two groups to determine whether they were statistically different.

### Mutant generation

The SARS-CoV-2 NTD construct with the native signal peptide was mutated by PCR mutagenesis to generate S13I and W152C mutations using the eponymously named primers (**Supplementary Table 1**). Plasmid sequences were verified by Genewiz sequencing facilities (Brooks Life Sciences).

Amino acid substitutions were introduced into the D614G pCDNA_SARS-CoV-2_S plasmid as previously described(*64*) using the QuikChange Lightening Site-Directed Mutagenesis kit, following the manufacturer’s instructions (Agilent Technologies, Inc., Santa Clara, CA). Sequences were checked by Sanger sequencing.

Plasmids encoding the SARS-CoV-2 S glycoprotein corresponding to the VOC B.1.427/B.1.429 (referred as B.1.429), B.1.1.7, B.1.351 and P.1 SARS-CoV-2 S glycoprotein-encoding-plasmids used to produce SARS-CoV-2-VSV, were obtained using a multistep based on overlap extension PCR (oePCR) protocol(*29*). Briefly, the mutations of the different VOC lineages were encoded on each primer pair used to amplify sequential, overlapping fragments of the SARS-CoV-2 D19 plasmid, which encodes a C-terminally truncated Spike proteins, known to support higher plasma membrane expression(*65*). Two to three contiguous PCR fragments were subsequently joined by oePCR using the most external primers. For all PCR reactions the Q5 Hot Start High fidelity DNA polymerase was used (New England Biolabs Inc.), according to the manufacturer’s instructions and adapting the elongation time to the size of the amplicon. After each PCR step the amplified regions were separated on agarose gel and purified using Illustra GFX™ PCR DNA and Gel Band Purification Kit (Merck KGaA). The last oePCR step was performed to amplify the complete SARS-CoV-2S D19 sequence using primers carrying 15bp long 5’ overhangs homologous to the vector backbone, the amplicon was then cloned into the pCDNA3 vector using the Takara In-fusion HD cloning kit, following manufacturer’s instructions.

### Production of California-B.1.429 (L452R) receptor binding domain and recombinant ectodomains

The SARS-CoV2 RBD-L452R construct California-B.1.429 (L452R) was synthesized by GenScript into CMVR with an N-terminal mu-phosphatase signal peptide and a C-terminal octa-histidine tag (GHHHHHHHH) and an avi tag. The boundaries of the construct are N-328RFPN331 and 528KKST531-C35. The B.1.429 RBD gene was synthesized by GenScript into pCMVR with the same boundaries and construct details with a mutation at L452R. This plasmid was transiently transfected into Expi293F cells using Expi293F expression medium (Life Technologies) at 37°C 8% CO2 rotating at 150 rpm. The culture was transfected using ExpiFectamine™ 293 Transfection Kit (Gibco, #A14524) and cultivated for 7 days. Supernatants were clarified by centrifugation (30 min at 4000xg) prior to loading onto a Strep-Tactin®XT 4Flow® high-capacity cartridge 5ml column (IBA-Lifesciences) and eluted with 50mM Biotin, 100 mM Tris-HCl, 150 mM NaCl, 1mM EDTA, pH 8.0, prior to buffer exchange by size exclusion chromatography (SEC) into formulation buffer 100mM Tris-HCl, 150 mM NaCl, 1mM EDTA, pH 8.0.

All SARS-CoV-2 S NTD domain constructs (residues 14-307) with a C-terminal 8XHis-tag were produced in 100 mL culture of Expi293F™ Cells (ThermoFisher Scientific) grown in suspension using Expi293™ Expression Medium (ThermoFisher Scientific) at 37°C in a humidified 8% CO2 incubator rotating at 130 r.p.m. Cells grown to a density of 3 million cells per mL were transfected using pCMV::SARS-CoV-2_S_NTD derivative mutants with the ExpiFectamine™ 293 Transfection Kit (ThermoFisher Scientific) with and cultivated for five days at which point the supernatant was harvested. His-tagged NTD domain constructs were purified from clarified supernatants using 2 ml of cobalt resin (Takara Bio TALON), washing with 50 column volumes of 20 mM HEPES-HCl pH 8.0 and 150 mM NaCl and eluted with 600 mM imidazole. Purified protein was concentrated using a 30 kDa centrifugal filter (Amicon Ultra 0.5 mL centrifugal filters, MilliporeSigma), the imidazole was washed away by consecutive dilutions in the centrifugal filter unit with 20 mM HEPES-HCl pH 8.0 and 150 mM NaCl, and finally concentrated to 20 mg/ml and flash frozen.

### Intact mass spectrometry analysis of purified NTD constructs

The purpose of intact MS was to verify the n-terminal sequence on the constructs. N-linked glycans were removed by PNGase F after overnight non-denaturing reaction at room temperature. 4 μg of deglycosylated protein was used for each injection on the LC-MS system to acquire intact MS signal after separation of protease and protein by LC (Agilent PLRP-S reversed phase column). Thermo MS (Q Exactive Plus Orbitrap) was used to acquire intact protein mass under denaturing condition. BioPharma Finder 3.2 software was used to deconvolute the raw m/z data to protein average mass.

### Non-reducing Peptide Mapping mass spectrometry analysis of purified NTD constructs

The purpose of peptide mapping was to verify the disulfide linkage between C136 and C152 on S13I/W152C variant. Combo protease with Glu-C and trypsin was used for protein digestion without adding reducing reagent. 50 μg of deglycosyated protein was denatured (6M guanidine hydrochloride), alkylated (Iodoacetamide), and buffer exchanged (Zeba spin desalting column) before digestion. 10 μg of digested peptide was analyzed on the LC-MS system (Agilent AdvanceBio peptide mapping column and Thermo Q Exactive Plus Orbitrap MS) to acquire both MS1 and MS2 data under HCD fragmentation. Peptide mapping data was analyzed on Biopharma Finder 3.2 by searching the possible disulfide linkages on the construct.

---

**Supplementary Table 1.**
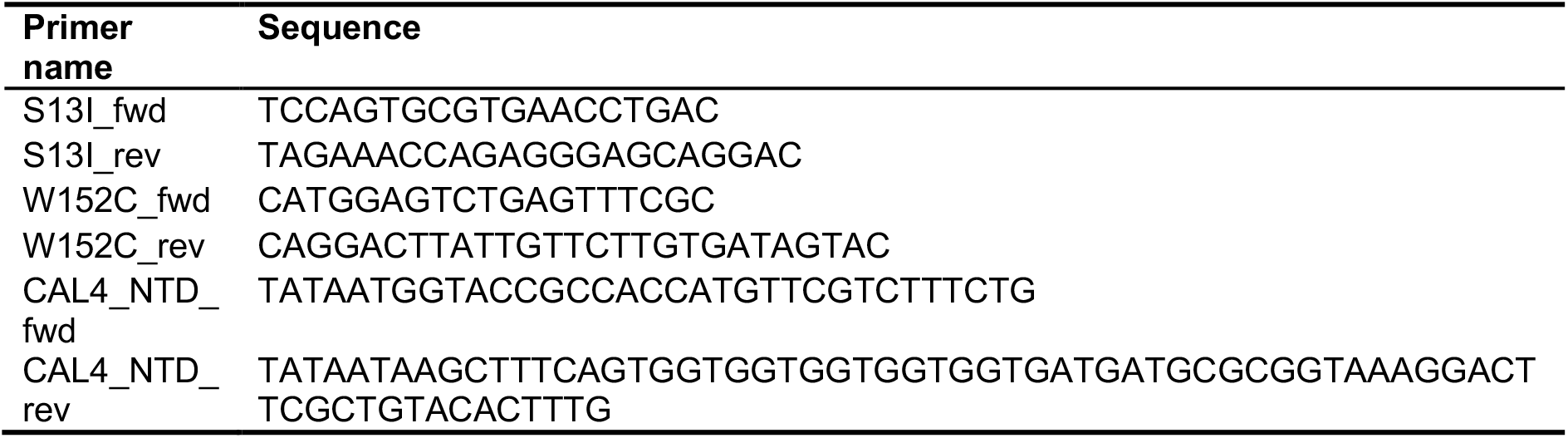
Primers used in this study.

**Supplemental Fig 1.**
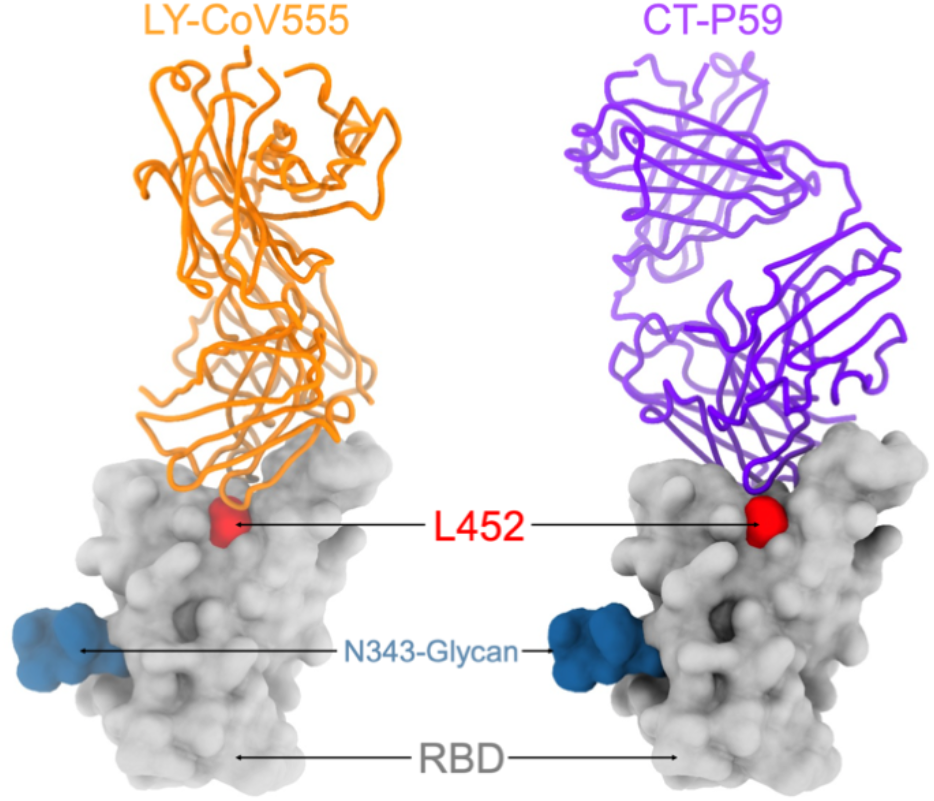
Surface representation of the SARS-CoV-2 RBD (grey) bound to the bamlanivimab (LY-CoV555, orange, PDB 7CM4) and regdanvimab (CT-P59, purple, PDB 7KMG) Fab fragments shown as ribbons. The L452 side chain is show as red spheres to indicate its central location within the epitopes of these two mAbs. The N343 glycan is rendered as blue spheres.

**Supplemental Fig. 2.**
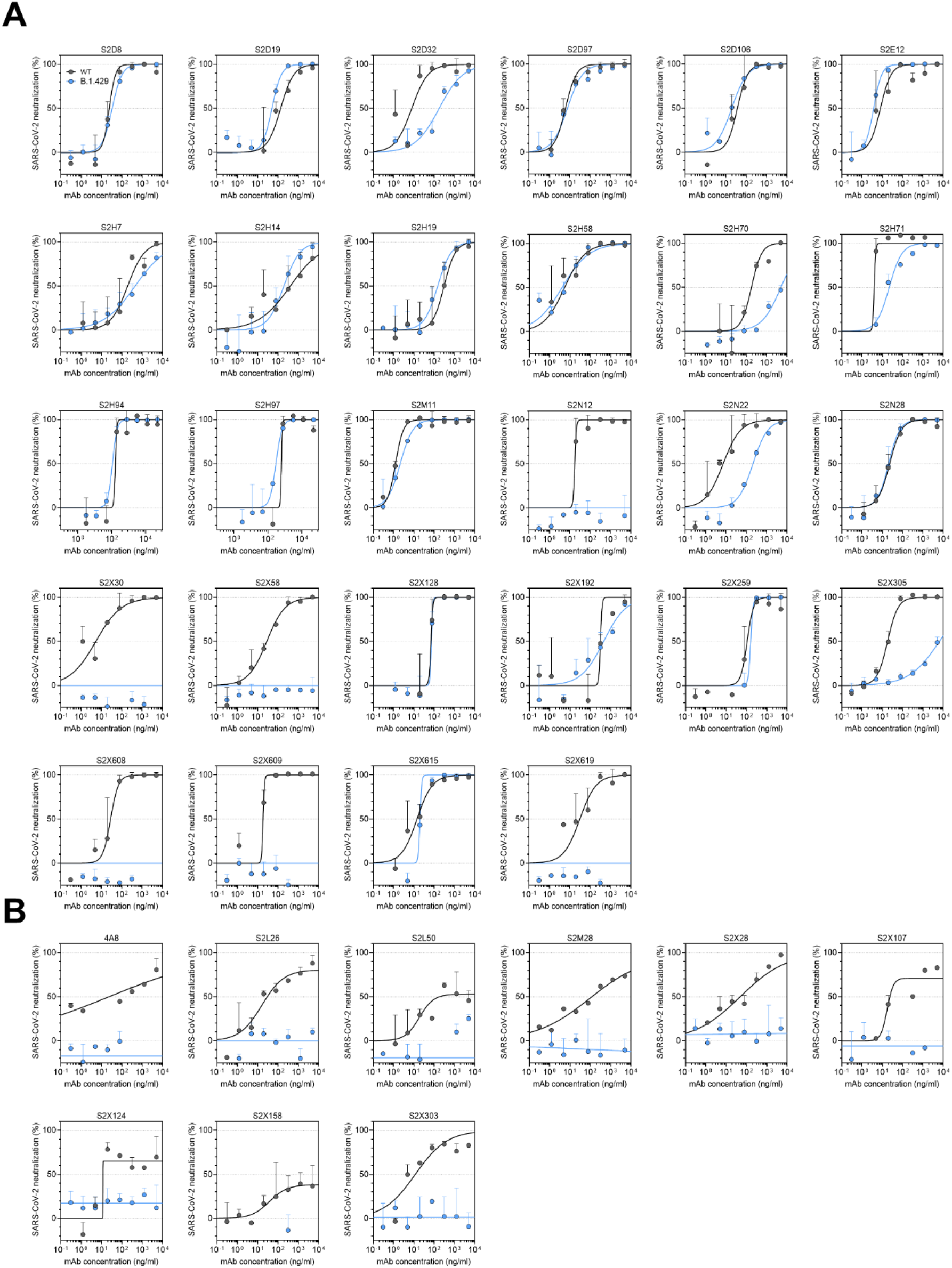
Neutralization by RBD- and NTD-specific mAbs against wildtype and B.1.427/B.1.429 SARS-CoV-2 S pseudoviruses. (**A,B**) Neutralization of SARS-CoV-2 pseudotyped VSV carrying wild-type D614 (grey) or B.1.427/B.1.429 (blue) S protein by RBD-targeting mAbs (A) and NTD-targeting mAbs (B). Data are representative of *n = 2* replicates.

**Supplemental Fig. 3.**
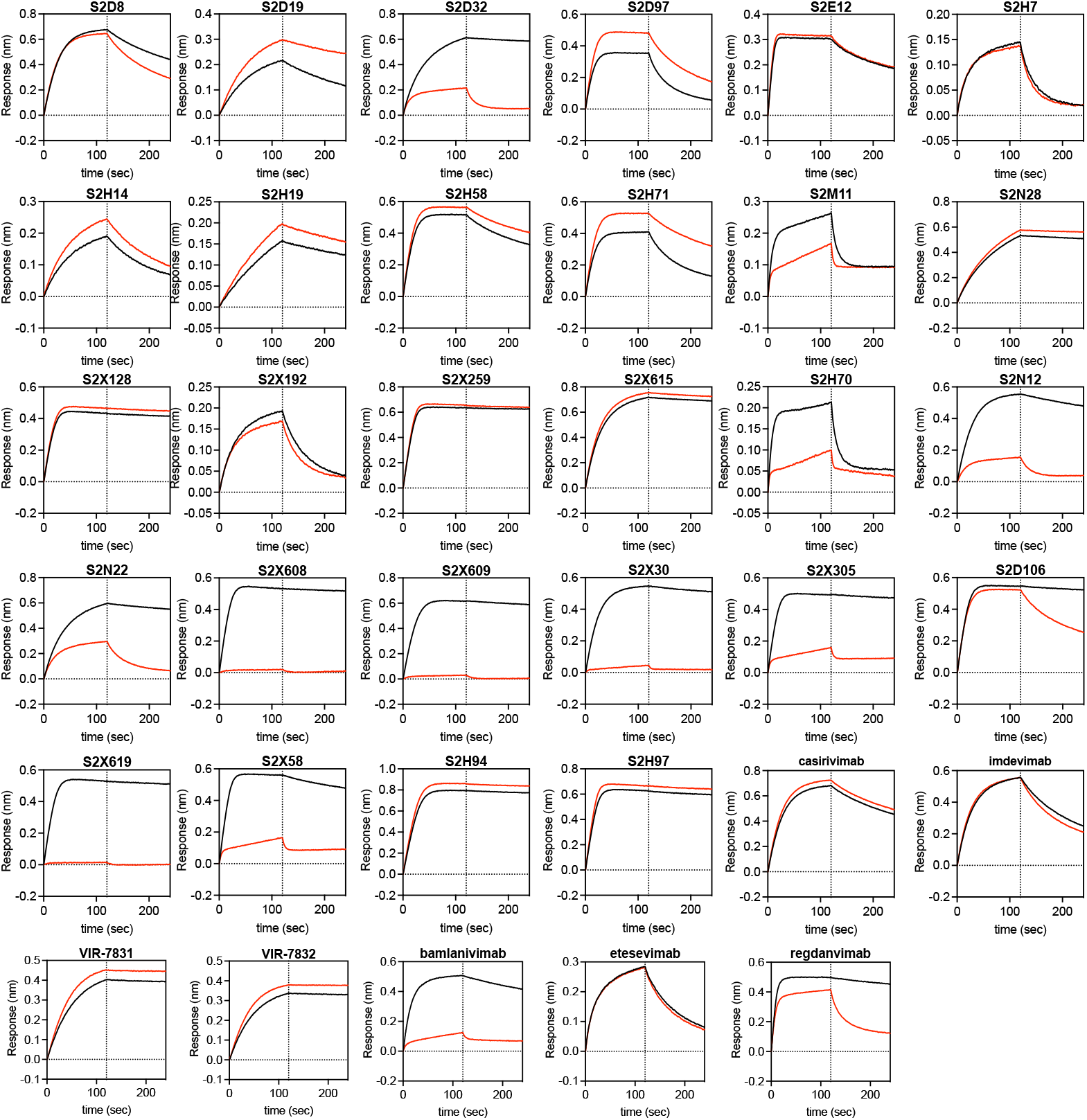
Kinetics of binding to wildtype and L452R SARS-CoV-2 RBD of 35 RBD-specific mAbs. Biolayer interferometry analysis of binding to wildtype (black) and L452R (red) RBDs by 35 RBD-targeting mAbs.

**Supplemental Fig. 4.**
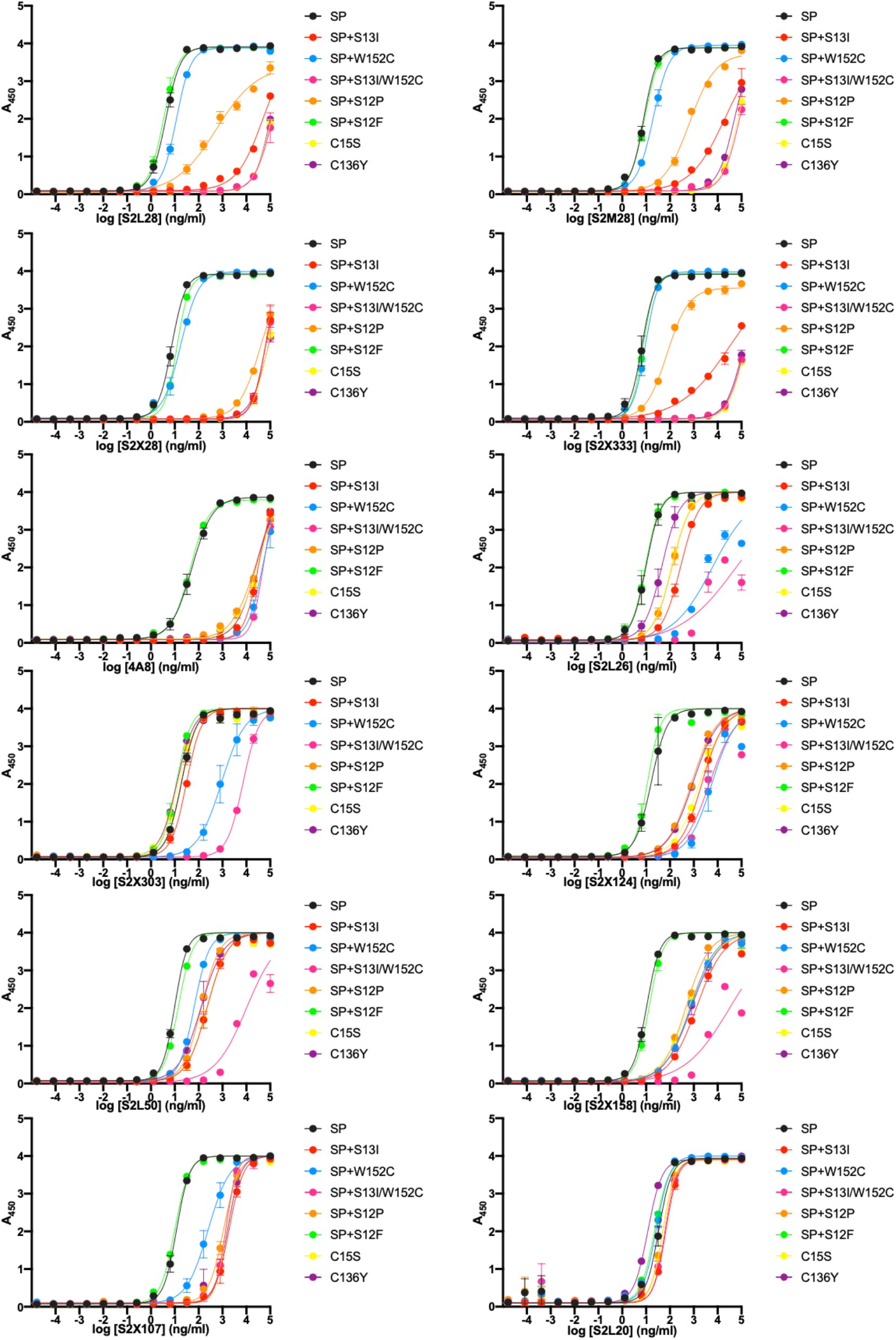
Effect of SARS-CoV-2 NTD mutations on mAb binding. Binding of a panel of 11 neutralizing (antigenic site i) and 1 non-neutralizing (antigenic site iv) NTD-specific mAbs to recombinant SARS-CoV-2 NTD variants analyzed by ELISA.

**Supplemental Fig. 5.**
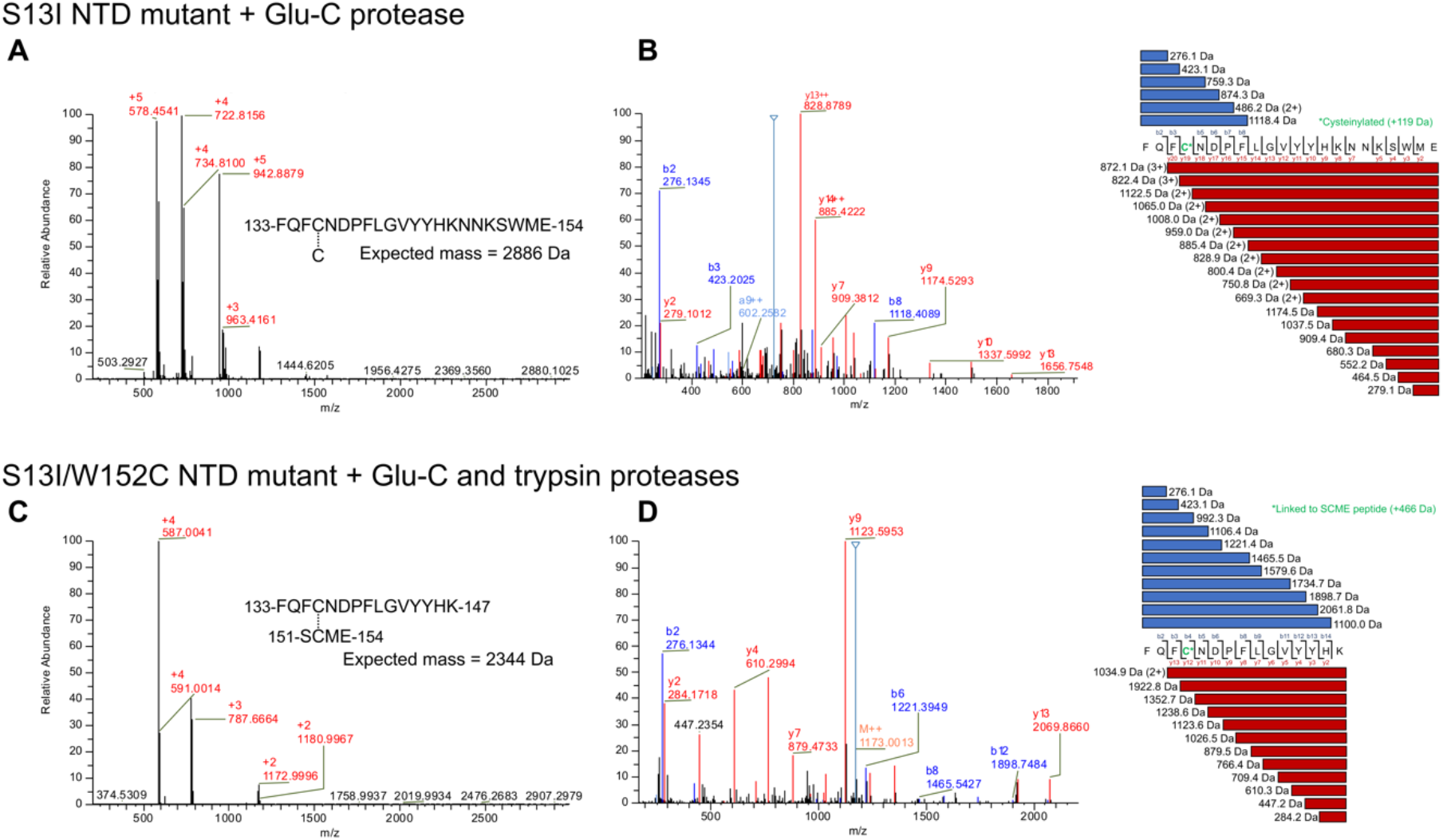
Mass spectrometry (MS) analysis of selected peptides. **A-B.** ESI-MS (A) and MS/MS (B) analysis of peptides containing cysteinylated residue C136 from the S13I NTD mutant treated with Glu-C protease. MS/MS analysis shows the MS plot with the most prominent peaks labelled (left) and a list of the identified fragmented peptides (right). **C-D.** ESI-MS (C) and MS/MS (D) analysis of peptides with a disulfide link between C136 and W152C from the S13I/W152C NTD mutant treated with Glu-C and trypsin proteases. MS/MS analysis shows the MS plot with the most prominent peaks labelled (left) and a list of the identified fragmented peptides (right).

## References

1. A. C. Walls, Y. J. Park, M. A. Tortorici, A. Wall, A. T. McGuire, D. Veesler, Structure, Function, and Antigenicity of the SARS-CoV-2 Spike Glycoprotein. Cell. 181, 281–292.e6 (2020).

2. D. Wrapp, N. Wang, K. S. Corbett, J. A. Goldsmith, C. L. Hsieh, O. Abiona, B. S. Graham, J. S. McLellan, Cryo-EM structure of the 2019-nCoV spike in the prefusion conformation. Science. 367, 1260–1263 (2020).

3. A. C. Walls, M. A. Tortorici, B. J. Bosch, B. Frenz, P. J. M. Rottier, F. DiMaio, F. A. Rey, D. Veesler, Cryo-electron microscopy structure of a coronavirus spike glycoprotein trimer. Nature. 531, 114–117 (2016).

4. M. A. Tortorici, D. Veesler, Structural insights into coronavirus entry. Adv. Virus Res. 105, 93–116 (2019).

5. M. Letko, A. Marzi, V. Munster, Functional assessment of cell entry and receptor usage for SARS-CoV-2 and other lineage B betacoronaviruses. Nature Microbiology (2020), doi:10.1038/s41564-020-0688-y.

6. P. Zhou, X. L. Yang, X. G. Wang, B. Hu, L. Zhang, W. Zhang, H. R. Si, Y. Zhu, B. Li, C. L. Huang, H. D. Chen, J. Chen, Y. Luo, H. Guo, R. D. Jiang, M. Q. Liu, Y. Chen, X. R. Shen, X. Wang, X. S. Zheng, K. Zhao, Q. J. Chen, F. Deng, L. L. Liu, B. Yan, F. X. Zhan, Y. Y. Wang, G. F. Xiao, Z. L. Shi, A pneumonia outbreak associated with a new coronavirus of probable bat origin. Nature (2020), doi:10.1038/s41586-020-2012-7.

7. M. Hoffmann, H. Kleine-Weber, S. Schroeder, N. Krüger, T. Herrler, S. Erichsen, T. S. Schiergens, G. Herrler, N. H. Wu, A. Nitsche, M. A. Müller, C. Drosten, S. Pöhlmann, SARS-CoV-2 Cell Entry Depends on ACE2 and TMPRSS2 and Is Blocked by a Clinically Proven Protease Inhibitor. Cell. 181, 271–280.e8 (2020).

8. W. T. Soh, Y. Liu, E. E. Nakayama, C. Ono, S. Torii, H. Nakagami, Y. Matsuura, T. Shioda, H. Arase, bioRxiv, in press.

9. S. Wang, Z. Qiu, Y. Hou, X. Deng, W. Xu, T. Zheng, P. Wu, S. Xie, W. Bian, C. Zhang, Z. Sun, K. Liu, C. Shan, A. Lin, S. Jiang, Y. Xie, Q. Zhou, L. Lu, J. Huang, X. Li, AXL is a candidate receptor for SARS-CoV-2 that promotes infection of pulmonary and bronchial epithelial cells. Cell Res. 31, 126–140 (2021).

10. D. Pinto, Y. J. Park, M. Beltramello, A. C. Walls, M. A. Tortorici, S. Bianchi, S. Jaconi, K. Culap, F. Zatta, A. De Marco, A. Peter, B. Guarino, R. Spreafico, E. Cameroni, J. B. Case, R. E. Chen, C. Havenar-Daughton, G. Snell, A. Telenti, H. W. Virgin, A. Lanzavecchia, M. Diamond, K. Fink, D. Veesler, D. Corti, Cross-neutralization of SARS-CoV-2 by a human monoclonal SARS-CoV antibody. Nature. 583, 290–295 (2020).

11. L. Piccoli, Y. J. Park, M. A. Tortorici, N. Czudnochowski, A. C. Walls, M. Beltramello, C. Silacci-Fregni, D. Pinto, L. E. Rosen, J. E. Bowen, O. J. Acton, S. Jaconi, B. Guarino, A. Minola, F. Zatta, N. Sprugasci, J. Bassi, A. Peter, A. De Marco, J. C. Nix, F. Mele, S. Jovic, B. F. Rodriguez, S. V. Gupta, F. Jin, G. Piumatti, G. Lo Presti, A. F. Pellanda, M. Biggiogero, M. Tarkowski, M. S. Pizzuto, E. Cameroni, C. Havenar-Daughton, M. Smithey, D. Hong, V. Lepori, E. Albanese, A. Ceschi, E. Bernasconi, L. Elzi, P. Ferrari, C. Garzoni, A. Riva, G. Snell, F. Sallusto, K. Fink, H. W. Virgin, A. Lanzavecchia, D. Corti, Veesler, Mapping Neutralizing and Immunodominant Sites on the SARS-CoV-2 Spike Receptor-Binding Domain by Structure-Guided High-Resolution Serology. Cell. 183, 1024–1042.e21 (2020).

12. M. McCallum, A. De Marco, F. A. Lempp, M. A. Tortorici, D. Pinto, A. C. Walls, M. Beltramello, A. Chen, Z. Liu, F. Zatta, S. Zepeda, J. di Iulio, J. E. Bowen, M. Montiel-Ruiz, J. Zhou, L. E. Rosen, S. Bianchi, B. Guarino, C. S. Fregni, R. Abdelnabi, S.-Y. Caroline Foo, P. W. Rothlauf, L.-M. Bloyet, F. Benigni, E. Cameroni, J. Neyts, A. Riva, G. Snell, A. Telenti, S. P. J. Whelan, H. W. Virgin, D. Corti, M. S. Pizzuto, D. Veesler, N-terminal domain antigenic mapping reveals a site of vulnerability for SARS-CoV-2. Cell (2021), doi:10.1016/j.cell.2021.03.028.

13. M. A. Tortorici, M. Beltramello, F. A. Lempp, D. Pinto, H. V. Dang, L. E. Rosen, M. McCallum, J. Bowen, A. Minola, S. Jaconi, F. Zatta, A. De Marco, B. Guarino, S. Bianchi, E. J. Lauron, H. Tucker, J. Zhou, A. Peter, C. Havenar-Daughton, J. A. Wojcechowskyj, J. B. Case, R. E. Chen, H. Kaiser, M. Montiel-Ruiz, M. Meury, N. Czudnochowski, R. Spreafico, J. Dillen, C. Ng, N. Sprugasci, K. Culap, F. Benigni, R. Abdelnabi, S. C. Foo, M. A. Schmid, E. Cameroni, A. Riva, A. Gabrieli, M. Galli, M. S. Pizzuto, J. Neyts, M. S. Diamond, H. W. Virgin, G. Snell, D. Corti, K. Fink, D. Veesler, Ultrapotent human antibodies protect against SARS-CoV-2 challenge via multiple mechanisms. Science. 370, 950–957 (2020).

14. J. Hansen, A. Baum, K. E. Pascal, V. Russo, S. Giordano, E. Wloga, B. O. Fulton, Y. Yan, K. Koon, K. Patel, K. M. Chung, A. Hermann, E. Ullman, J. Cruz, A. Rafique, T. Huang, J. Fairhurst, C. Libertiny, M. Malbec, W. Y. Lee, R. Welsh, G. Farr, S. Pennington, D. Deshpande, J. Cheng, A. Watty, P. Bouffard, R. Babb, N. Levenkova, C. Chen, B. Zhang, A. Romero Hernandez, K. Saotome, Y. Zhou, M. Franklin, S. Sivapalasingam, D. C. Lye, S. Weston, J. Logue, R. Haupt, M. Frieman, G. Chen, W. Olson, A. J. Murphy, N. Stahl, G. D. Yancopoulos, C. A. Kyratsous, Studies in humanized mice and convalescent humans yield a SARS-CoV-2 antibody cocktail. Science (2020), doi:10.1126/science.abd0827.

15. A. Baum, B. O. Fulton, E. Wloga, R. Copin, K. E. Pascal, V. Russo, S. Giordano, K. Lanza, N. Negron, M. Ni, Y. Wei, G. S. Atwal, A. J. Murphy, N. Stahl, G. D. Yancopoulos, C. A. Kyratsous, Antibody cocktail to SARS-CoV-2 spike protein prevents rapid mutational escape seen with individual antibodies. Science (2020), doi:10.1126/science.abd0831.

16. N. Suryadevara, S. Shrihari, P. Gilchuk, L. A. VanBlargan, E. Binshtein, S. J. Zost, R. S. Nargi, R. E. Sutton, E. S. Winkler, E. C. Chen, M. E. Fouch, E. Davidson, B. J. Doranz, R. E. Chen, P.-Y. Shi, R. H. Carnahan, L. B. Thackray, M. S. Diamond, J. E. Crowe Jr, Neutralizing and protective human monoclonal antibodies recognizing the N-terminal domain of the SARS-CoV-2 spike protein. Cell (2021), doi:10.1016/j.cell.2021.03.029.

17. G. Cerutti, Y. Guo, T. Zhou, J. Gorman, M. Lee, M. Rapp, E. R. Reddem, J. Yu, F. Bahna, J. Bimela, Y. Huang, P. S. Katsamba, L. Liu, M. S. Nair, R. Rawi, A. S. Olia, P. Wang, B. Zhang, G.-Y. Chuang, D. D. Ho, Z. Sheng, P. D. Kwong, L. Shapiro, Potent SARS-CoV-2 neutralizing antibodies directed against spike N-terminal domain target a single supersite. Cell Host Microbe (2021), doi:10.1016/j.chom.2021.03.005.

18. X. Chi, R. Yan, J. Zhang, G. Zhang, Y. Zhang, M. Hao, Z. Zhang, P. Fan, Y. Dong, Y. Yang, Z. Chen, Y. Guo, Y. Li, X. Song, Y. Chen, L. Xia, L. Fu, L. Hou, J. Xu, C. Yu, J. Li, Q. Zhou, W. Chen, A neutralizing human antibody binds to the N-terminal domain of the Spike protein of SARS-CoV-2. Science. 369, 650–655 (2020).

19. Z. Wang, F. Schmidt, Y. Weisblum, F. Muecksch, C. O. Barnes, S. Finkin, D. Schaefer-Babajew, M. Cipolla, C. Gaebler, J. A. Lieberman, T. Y. Oliveira, Z. Yang, M. E. Abernathy, K. E. Huey-Tubman, A. Hurley, M. Turroja, K. A. West, K. Gordon, K. G. Millard, V. Ramos, J. D. Silva, J. Xu, R. A. Colbert, R. Patel, J. Dizon, C. Unson-O’Brien, I. Shimeliovich, A. Gazumyan, M. Caskey, P. J. Bjorkman, R. Casellas, T. Hatziioannou, P. D. Bieniasz, M. C. Nussenzweig, mRNA vaccine-elicited antibodies to SARS-CoV-2 and circulating variants. Nature (2021), doi:10.1038/s41586-021-03324-6.

20. C. O. Barnes, C. A. Jette, M. E. Abernathy, K.-M. A. Dam, S. R. Esswein, H. B. Gristick, A. G. Malyutin, N. G. Sharaf, K. E. Huey-Tubman, Y. E. Lee, D. F. Robbiani, M. C. Nussenzweig, A. P. West Jr, P. J. Bjorkman, SARS-CoV-2 neutralizing antibody structures inform therapeutic strategies. Nature. 588, 682–687 (2020).

21. D. F. Robbiani, C. Gaebler, F. Muecksch, J. C. C. Lorenzi, Z. Wang, A. Cho, M. Agudelo, C. O. Barnes, A. Gazumyan, S. Finkin, T. Hägglöf, T. Y. Oliveira, C. Viant, A. Hurley, H. H. Hoffmann, K. G. Millard, R. G. Kost, M. Cipolla, K. Gordon, F. Bianchini, S. T. Chen, V. Ramos, R. Patel, J. Dizon, I. Shimeliovich, P. Mendoza, H. Hartweger, L. Nogueira, M. Pack, J. Horowitz, F. Schmidt, Y. Weisblum, E. Michailidis, A. W. Ashbrook, E. Waltari, J. E. Pak, K. E. Huey-Tubman, N. Koranda, P. R. Hoffman, A. P. West, C. M. Rice, T. Hatziioannou, P. J. Bjorkman, P. D. Bieniasz, M. Caskey, M. C. Nussenzweig, Convergent antibody responses to SARS-CoV-2 in convalescent individuals. Nature (2020), doi:10.1038/s41586-020-2456-9.

22. B. E. Jones, P. L. Brown-Augsburger, K. S. Corbett, K. Westendorf, J. Davies, T. P. Cujec, C. M. Wiethoff, J. L. Blackbourne, B. A. Heinz, D. Foster, R. E. Higgs, D. Balasubramaniam, L. Wang, R. Bidshahri, L. Kraft, Y. Hwang, S. Žentelis, K. R. Jepson, R. Goya, M. A. Smith, D. W. Collins, S. J. Hinshaw, S. A. Tycho, D. Pellacani, P. Xiang, K. Muthuraman, S. Sobhanifar, M. H. Piper, F. J. Triana, J. Hendle, A. Pustilnik, A. C. Adams, S. J. Berens, R. S. Baric, D. R. Martinez, R. W. Cross, T. W. Geisbert, V. Borisevich, O. Abiona, H. M. Belli, M. de Vries, A. Mohamed, M. Dittmann, M. Samanovic, M. J. Mulligan, J. A. Goldsmith, C. L. Hsieh, N. V. Johnson, D. Wrapp, J. S. McLellan, B. C. Barnhart, B. S. Graham, J. R. Mascola, C. L. Hansen, E. Falconer, LY-CoV555, a rapidly isolated potent neutralizing antibody, provides protection in a non-human primate model of SARS-CoV-2 infection. bioRxiv (2020), doi:10.1101/2020.09.30.318972.

23. M. M. Sauer, M. A. Tortorici, Y.-J. Park, A. C. Walls, L. Homad, O. Acton, J. Bowen, C. Wang, X. Xiong, W. de van der Schueren, J. Quispe, B. G. Hoffstrom, B.-J. Bosch, A. T. McGuire, D. Veesler, Structural basis for broad coronavirus neutralization. bioRxiv (2020), doi:10.1101/2020.12.29.424482.

24. C. Wang, R. van Haperen, J. Gutiérrez-Álvarez, W. Li, N. M. A. Okba, I. Albulescu, I. Widjaja, B. van Dieren, R. Fernandez-Delgado, I. Sola, D. L. Hurdiss, O. Daramola, F. Grosveld, F. J. M. van Kuppeveld, B. L. Haagmans, L. Enjuanes, D. Drabek, B.-J. Bosch, A conserved immunogenic and vulnerable site on the coronavirus spike protein delineated by cross-reactive monoclonal antibodies. Nat. Commun. 12, 1715 (2021).

25. G. Song, W.-T. He, S. Callaghan, F. Anzanello, D. Huang, J. Ricketts, J. L. Torres, N. Beutler, L. Peng, S. Vargas, J. Cassell, M. Parren, L. Yang, C. Ignacio, D. M. Smith, J. E. Voss, D. Nemazee, A. B. Ward, T. Rogers, D. R. Burton, R. Andrabi, bioRxiv, in press.

26. H. Tegally, E. Wilkinson, M. Giovanetti, A. Iranzadeh, V. Fonseca, J. Giandhari, D. Doolabh, S. Pillay, E. J. San, N. Msomi, K. Mlisana, A. von Gottberg, S. Walaza, M. Allam, A. Ismail, T. Mohale, A. J. Glass, S. Engelbrecht, G. Van Zyl, W. Preiser, F. Petruccione, A. Sigal, D. Hardie, G. Marais, M. Hsiao, S. Korsman, M.-A. Davies, L. Tyers, I. Mudau, D. York, C. Maslo, D. Goedhals, S. Abrahams, O. Laguda-Akingba, A. Alisoltani-Dehkordi, A. Godzik, C. K. Wibmer, B. T. Sewell, J. Lourenço, L. C. J. Alcantara, S. L. Kosakovsky Pond, S. Weaver, D. Martin, R. J. Lessells, J. N. Bhiman, C. Williamson, T. de Oliveira, Emergence of a SARS-CoV-2 variant of concern with mutations in spike glycoprotein. Nature (2021), doi:10.1038/s41586-021-03402-9.

27. N. R. Faria, I. M. Claro, D. Candido, L. A. Moyses Franco, P. S. Andrade, T. M. Coletti, C. A. M. Silva, F. C. Sales, E. R. Manuli, R. S. Aguiar, Others, Genomic characterisation of an emergent SARS-CoV-2 lineage in Manaus: preliminary findings. January. 12, 2021 (2021).

28. N. G. Davies, S. Abbott, R. C. Barnard, C. I. Jarvis, A. J. Kucharski, J. D. Munday, C. A. B. Pearson, T. W. Russell, D. C. Tully, A. D. Washburne, T. Wenseleers, A. Gimma, W. Waites, K. L. M. Wong, K. van Zandvoort, J. D. Silverman, CMMID COVID-19 Working Group, COVID-19 Genomics UK (COG-UK) Consortium, K. Diaz-Ordaz, R. Keogh, R. M. Eggo, S. Funk, M. Jit, K. E. Atkins, W. J. Edmunds, Estimated transmissibility and impact of SARS-CoV-2 lineage B.1.1.7 in England. Science (2021), doi:10.1126/science.abg3055.

29. D. A. Collier, A. De Marco, I. A. T. M. Ferreira, B. Meng, R. Datir, A. C. Walls, S. A. Kemp S, J. Bassi, D. Pinto, C. S. Fregni, S. Bianchi, M. A. Tortorici, J. Bowen, K. Culap, S. Jaconi, E. Cameroni, G. Snell, M. S. Pizzuto, A. F. Pellanda, C. Garzoni, A. Riva, A. Elmer, N. Kingston, B. Graves, L. E. McCoy, K. G. C. Smith, J. R. Bradley, N. Temperton, L. Lourdes Ceron-Gutierrez, G. Barcenas-Morales, W. Harvey, H. W. Virgin, A. Lanzavecchia, L. Piccoli, R. Doffinger, M. Wills, D. Veesler, D. Corti, R. K. Gupta, The CITIID-NIHR BioResource COVID-19 Collaboration, The COVID-19 Genomics UK (COG-UK) consortium, Sensitivity of SARS-CoV-2 B.1.1.7 to mRNA vaccine-elicited antibodies. Nature (2021), doi:10.1038/s41586-021-03412-7.

30. P. Wang, M. S. Nair, L. Liu, S. Iketani, Y. Luo, Y. Guo, M. Wang, J. Yu, B. Zhang, P. D. Kwong, B. S. Graham, J. R. Mascola, J. Y. Chang, M. T. Yin, M. Sobieszczyk, C. A. Kyratsous, L. Shapiro, Z. Sheng, Y. Huang, D. D. Ho, Antibody resistance of SARS-CoV-2 variants B.1.351 and B.1.1.7. Nature (2021), doi:10.1038/s41586-021-03398-2.

31. X. Deng, M. A. Garcia-Knight, M. M. Khalid, V. Servellita, C. Wang, M. K. Morris, A. Sotomayor-Gonzalez, D. R. Glasner, K. R. Reyes, A. S. Gliwa, N. P. Reddy, C. Sanchez San Martin, S. Federman, J. Cheng, J. Balcerek, J. Taylor, J. A. Streithorst, S. Miller, G. R. Kumar, B. Sreekumar, P.-Y. Chen, U. Schulze-Gahmen, T. Y. Taha, J. M. Hayashi, C. R. Siomoneau, S. McMahon, P. V. Lidsky, Y. Xiao, N. M. Green, P. Hemarajata, A. Espinosa, C. Kath, M. Haw, J. Bell, J. K. Hacker, C. Hanson, D. A. Wadford, C. Anaya, D. Ferguson, L. F. Lareau, P. A. Frankino, H. Shivram, S. K. Wyman, M. Ott, R. Andino, C. Y. Chiu, Transmission, infectivity, and antibody neutralization of an emerging SARS-CoV-2 variant in California carrying a L452R spike protein mutation. bioRxiv (2021),, doi:10.1101/2021.03.07.21252647.

32. A. Rambaut, E. C. Holmes, Á. O’Toole, V. Hill, J. T. McCrone, C. Ruis, L. du Plessis, O. G. Pybus, A dynamic nomenclature proposal for SARS-CoV-2 lineages to assist genomic epidemiology. Nat. Microbiol. 5, 1403–1407 (2020).

33. W. Zhang, B. D. Davis, S. S. Chen, J. M. Sincuir Martinez, J. T. Plummer, E. Vail, Emergence of a novel SARS-CoV-2 variant in southern California. JAMA (2021), doi:10.1001/jama.2021.1612.

34. V. Tchesnokova, H. Kulakesara, L. Larson, V. Bowers, E. Rechkina, D. Kisiela, Y. Sledneva, D. Choudhury, I. Maslova, K. Deng, K. Kutumbaka, H. Geng, C. Fowler, D. Greene, J. Ralston, M. Samadpour, E. Sokurenko, Acquisition of the L452R mutation in the ACE2-binding interface of Spike protein triggers recent massive expansion of SARS-Cov-2 variants. bioRxivorg (2021), doi:10.1101/2021.02.22.432189.

35. T. Moyo-Gwete, M. Madzivhandila, Z. Makhado, F. Ayres, D. Mhlanga, B. Oosthuysen, B. E. Lambson, P. Kgagudi, H. Tegally, A. Iranzadeh, D. Doolabh, L. Tyers, L. R. Chinhoyi, M. Mennen, S. Skelm, C. K. Wibmer, J. N. Bhiman, V. Ueckermann, T. Rossouw, M. Boswell, T. de Oliveira, C. Williamson, W. A. Burgers, N. Ntusi, L. Morris, P. L. Moore, SARS-CoV-2 501Y.V2 (B.1.351) elicits cross-reactive neutralizing antibodies. bioRxivorg (2021), doi:10.1101/2021.03.06.434193.

36. S. Cele, I. Gazy, L. Jackson, S.-H. Hwa, H. Tegally, G. Lustig, J. Giandhari, S. Pillay, E. Wilkinson, Y. Naidoo, F. Karim, Y. Ganga, K. Khan, M. Bernstein, A. B. Balazs, B. I. Gosnell, W. Hanekom, M.-Y. S. Moosa, NGS-SA, COMMIT-KZN Team, R. J. Lessells, T. de Oliveira, A. Sigal, Escape of SARS-CoV-2 501Y.V2 from neutralization by convalescent plasma. Nature (2021), doi:10.1038/s41586-021-03471-w.

37. J. K. Millet, G. R. Whittaker, Murine Leukemia Virus (MLV)-based Coronavirus Spike-pseudotyped Particle Production and Infection. Bio Protoc. 6 (2016), doi:10.21769/BioProtoc.2035.

38. A. L. Cathcart, C. Havenar-Daughton, F. A. Lempp, D. Ma, M. Schmid, M. L. Agostini, B. Guarino, J. Di iulio, L. Rosen, H. Tucker, J. Dillen, S. Subramanian, B. Sloan, S. Bianchi, J. Wojcechowskyj, J. Zhou, H. Kaiser, A. Chase, M. Montiel-Ruiz, N. Czudnochowski, E. Cameroni, S. Ledoux, C. Colas, L. Soriaga, A. Telenti, S. Hwang, G. Snell, H. W. Virgin, D. Corti, C. M. Hebner, The dual function monoclonal antibodies VIR-7831 and VIR-7832 demonstrate potent in vitro and in vivo activity against SARS-CoV-2. bioRxiv (2021),, doi:10.1101/2021.03.09.434607.

39. P. S Arunachalam, A. C. Walls, N. Golden, C. Atyeo, S. Fischinger, C. Li, P. Aye, M. J. Navarro, L. Lai, V. V. Edara, K. Roltgen, K. Rogers, L. Shirreff, D. E. Ferrell, S. Wrenn, D. Pettie, J. C. Kraft, M. C. Miranda, E. Kepl, C. Sydeman, N. Brunette, M. Murphy, B. Fiala, L. Carter, A. G. White, M. Trisal, C.-L. Hsieh, K. Russell-Lodrigue, C. Monjure, J. Dufour, L. Doyle-Meyer, R. B. Bohm, N. J. Maness, C. Roy, J. A. Plante, K. S. Plante, A. Zhu, M. J. Gorman, S. Shin, X. Shen, J. Fontenot, S. Gupta, D. T. O Hagan, R. V. D. Most, R. Rappuoli, R. L. Coffman, D. Novack, J. S. McLellan, S. Subramaniam, D. Montefiori, S. D. Boyd, J. L. Flynn, G. Alter, F. Villinger, H. Kleanthous, J. Rappaport, M. Suthar, N. P. King, D. Veesler, B. Pulendran, Adjuvanting a subunit SARS-CoV-2 nanoparticle vaccine to induce protective immunity in non-human primates. bioRxiv (2021), doi:10.1101/2021.02.10.430696.

40. K. McMahan, J. Yu, N. B. Mercado, C. Loos, L. H. Tostanoski, A. Chandrashekar, J. Liu, L. Peter, C. Atyeo, A. Zhu, E. A. Bondzie, G. Dagotto, M. S. Gebre, C. Jacob-Dolan, Z. Li, F. Nampanya, S. Patel, L. Pessaint, A. Van Ry, K. Blade, J. Yalley-Ogunro, M. Cabus, R. Brown, A. Cook, E. Teow, H. Andersen, M. G. Lewis, D. A. Lauffenburger, G. Alter, D. H. Barouch, Correlates of protection against SARS-CoV-2 in rhesus macaques. Nature. 590, 630–634 (2021).

41. A. Baum, D. Ajithdoss, R. Copin, A. Zhou, K. Lanza, N. Negron, M. Ni, Y. Wei, K. Mohammadi, B. Musser, G. S. Atwal, A. Oyejide, Y. Goez-Gazi, J. Dutton, E. Clemmons, H. M. Staples, C. Bartley, B. Klaffke, K. Alfson, M. Gazi, O. Gonzalez, E. Dick, R. Carrion, L. Pessaint, M. Porto, A. Cook, R. Brown, V. Ali, J. Greenhouse, T. Taylor, H. Andersen, G. Lewis, N. Stahl, A. J. Murphy, G. D. Yancopoulos, C. A. Kyratsous, REGN-COV2 antibodies prevent and treat SARS-CoV-2 infection in rhesus macaques and hamsters. Science (2020), doi:10.1126/science.abe2402.

42. C. K. Wibmer, F. Ayres, T. Hermanus, M. Madzivhandila, P. Kgagudi, B. Oosthuysen, B. E. Lambson, T. de Oliveira, M. Vermeulen, K. van der Berg, T. Rossouw, M. Boswell, V. Ueckermann, S. Meiring, A. von Gottberg, C. Cohen, L. Morris, J. N. Bhiman, P. L. Moore, SARS-CoV-2 501Y.V2 escapes neutralization by South African COVID-19 donor plasma. Nat. Med. (2021), doi:10.1038/s41591-021-01285-x.

43. W. F. Garcia-Beltran, E. C. Lam, K. St Denis, A. D. Nitido, Z. H. Garcia, B. M. Hauser, J. Feldman, M. N. Pavlovic, D. J. Gregory, M. C. Poznansky, A. Sigal, A. G. Schmidt, A. J. Iafrate, V. Naranbhai, A. B. Balazs, Multiple SARS-CoV-2 variants escape neutralization by vaccine-induced humoral immunity. Cell (2021), doi:10.1016/j.cell.2021.03.013.

44. V. V. Edara, C. Norwood, K. Floyd, L. Lai, M. E. Davis-Gardner, W. H. Hudson, G. Mantus, L. E. Nyhoff, M. W. Adelman, R. Fineman, S. Patel, R. Byram, D. N. Gomes, G. Michael, H. Abdullahi, N. Beydoun, B. Panganiban, N. McNair, K. Hellmeister, J. Pitts, J. Winters, J. Kleinhenz, J. Usher, J. B. O’Keefe, A. Piantadosi, J. J. Waggoner, A. Babiker, D. S. Stephens, E. J. Anderson, S. Edupuganti, N. Rouphael, R. Ahmed, J. Wrammert, M. S. Suthar, Infection and vaccine-induced antibody binding and neutralization of the B.1.351 SARS-CoV-2 variant. Cell Host Microbe (2021), doi:10.1016/j.chom.2021.03.009.

45. Z. Liu, L. A. VanBlargan, L.-M. Bloyet, P. W. Rothlauf, R. E. Chen, S. Stumpf, H. Zhao, J. M. Errico, E. S. Theel, M. J. Liebeskind, B. Alford, W. J. Buchser, A. H. Ellebedy, D. H. Fremont, M. S. Diamond, S. P. J. Whelan, Identification of SARS-CoV-2 spike mutations that attenuate monoclonal and serum antibody neutralization. Cell Host Microbe (2021), doi:10.1016/j.chom.2021.01.014.

46. Q. Li, J. Wu, J. Nie, L. Zhang, H. Hao, S. Liu, C. Zhao, Q. Zhang, H. Liu, L. Nie, H. Qin, M. Wang, Q. Lu, X. Li, Q. Sun, J. Liu, W. Huang, Y. Wang, The Impact of Mutations in SARS-CoV-2 Spike on Viral Infectivity and Antigenicity. Cell. 182, 1284–1294.e9 (2020).

47. T. N. Starr, A. J. Greaney, A. S. Dingens, J. D. Bloom, Complete map of SARS-CoV-2 RBD mutations that escape the monoclonal antibody LY-CoV555 and its cocktail with LY-CoV016,, doi:10.1101/2021.02.17.431683.

48. T. N. Starr, A. J. Greaney, S. K. Hilton, D. Ellis, K. H. D. Crawford, A. S. Dingens, M. J. Navarro, J. E. Bowen, M. A. Tortorici, A. C. Walls, N. P. King, D. Veesler, J. D. Bloom, Deep Mutational Scanning of SARS-CoV-2 Receptor Binding Domain Reveals Constraints on Folding and ACE2 Binding. Cell. 182, 1295–1310.e20 (2020).

49. E. C. Thomson, L. E. Rosen, J. G. Shepherd, R. Spreafico, A. da Silva Filipe, J. A. Wojcechowskyj, C. Davis, L. Piccoli, D. J. Pascall, J. Dillen, S. Lytras, N. Czudnochowski, R. Shah, M. Meury, N. Jesudason, A. De Marco, K. Li, J. Bassi, A. O’Toole, D. Pinto, R. M. Colquhoun, K. Culap, B. Jackson, F. Zatta, A. Rambaut, S. Jaconi, V. B. Sreenu, J. Nix, I. Zhang, R. F. Jarrett, W. G. Glass, M. Beltramello, K. Nomikou, M. Pizzuto, L. Tong, E. Cameroni, T. I. Croll, N. Johnson, J. Di Iulio, A. Wickenhagen, A. Ceschi, A. M. Harbison, D. Mair, P. Ferrari, K. Smollett, F. Sallusto, S. Carmichael, C. Garzoni, J. Nichols, M. Galli, J. Hughes, A. Riva, A. Ho, M. Schiuma, M. G. Semple, P. J. M. Openshaw, E. Fadda, J. K. Baillie, J. D. Chodera, S. J. Rihn, S. J. Lycett, H. W. Virgin, A. Telenti, D. Corti, D. L. Robertson, G. Snell, Circulating SARS-CoV-2 spike N439K variants maintain fitness while evading antibody-mediated immunity. Cell (2021), doi:10.1016/j.cell.2021.01.037.

50. V. A. Avanzato, M. J. Matson, S. N. Seifert, R. Pryce, B. N. Williamson, S. L. Anzick, K. Barbian, S. D. Judson, E. R. Fischer, C. Martens, T. A. Bowden, E. de Wit, F. X. Riedo, V. J. Munster, Case Study: Prolonged infectious SARS-CoV-2 shedding from an asymptomatic immunocompromised cancer patient. Cell (2020), doi:10.1016/j.cell.2020.10.049.

51. B. Choi, M. C. Choudhary, J. Regan, J. A. Sparks, R. F. Padera, X. Qiu, I. H. Solomon, H. H. Kuo, J. Boucau, K. Bowman, U. D. Adhikari, M. L. Winkler, A. A. Mueller, T. Y. Hsu, M. Desjardins, L. R. Baden, B. T. Chan, B. D. Walker, M. Lichterfeld, M. Brigl, D. S. Kwon, S. Kanjilal, E. T. Richardson, A. H. Jonsson, G. Alter, A. K. Barczak, W. P. Hanage, X. G. Yu, G. D. Gaiha, M. S. Seaman, M. Cernadas, J. Z. Li, Persistence and Evolution of SARS-CoV-2 in an Immunocompromised Host. N. Engl. J. Med. (2020), doi:10.1056/NEJMc2031364.

52. K. R. McCarthy, L. J. Rennick, S. Nambulli, L. R. Robinson-McCarthy, W. G. Bain, G. Haidar, W. P. Duprex, Recurrent deletions in the SARS-CoV-2 spike glycoprotein drive antibody escape. Science (2021), doi:10.1126/science.abf6950.

53. E. Andreano, G. Piccini, D. Licastro, L. Casalino, N. V. Johnson, I. Paciello, S. D. Monego, E. Pantano, N. Manganaro, A. Manenti, R. Manna, E. Casa, I. Hyseni, L. Benincasa, E. Montomoli, R. E. Amaro, J. S. McLellan, R. Rappuoli, bioRxiv, in press.

54. Y. Weisblum, F. Schmidt, F. Zhang, J. DaSilva, D. Poston, J. C. C. Lorenzi, F. Muecksch, M. Rutkowska, H.-H. Hoffmann, E. Michailidis, C. Gaebler, M. Agudelo, A. Cho, Z. Wang, A. Gazumyan, M. Cipolla, L. Luchsinger, C. D. Hillyer, M. Caskey, D. F. Robbiani, C. M. Rice, M. C. Nussenzweig, T. Hatziioannou, P. D. Bieniasz, Escape from neutralizing antibodies by SARS-CoV-2 spike protein variants. Elife. 9, e61312 (2020).

55. M. K. Annavajhala, H. Mohri, J. E. Zucker, Z. Sheng, P. Wang, A. Gomez-Simmonds, D. D. Ho, A.-C. Uhlemann, A novel SARS-CoV-2 variant of concern, B.1.526, identified in New York. medRxiv (2021), doi:10.1101/2021.02.23.21252259.

56. A. P. West Jr, C. O. Barnes, Z. Yang, P. J. Bjorkman, SARS-CoV-2 lineage B.1.526 emerging in the New York region detected by software utility created to query the spike mutational landscape. bioRxiv (2021),, doi:10.1101/2021.02.14.431043.

57. G. S. C. Slater, E. Birney, Automated generation of heuristics for biological sequence comparison. BMC Bioinformatics. 6, 31 (2005).

58. K. Katoh, D. M. Standley, MAFFT multiple sequence alignment software version 7: improvements in performance and usability. Mol. Biol. Evol. 30, 772–780 (2013).

59. D. Corti, J. Voss, S. J. Gamblin, G. Codoni, A. Macagno, D. Jarrossay, S. G. Vachieri, D. Pinna, A. Minola, F. Vanzetta, C. Silacci, B. M. Fernandez-Rodriguez, G. Agatic, S. Bianchi, I. Giacchetto-Sasselli, L. Calder, F. Sallusto, P. Collins, L. F. Haire, N. Temperton, J. P. Langedijk, J. J. Skehel, A. Lanzavecchia, A neutralizing antibody selected from plasma cells that binds to group 1 and group 2 influenza A hemagglutinins. Science. 333, 850–856 (2011).

60. A. Takada, C. Robison, H. Goto, A. Sanchez, K. G. Murti, M. A. Whitt, Y. Kawaoka, A system for functional analysis of Ebola virus glycoprotein. Proc. Natl. Acad. Sci. U. S. A. 94, 14764–14769 (1997).

61. A. M. Riblett, V. A. Blomen, L. T. Jae, L. A. Altamura, R. W. Doms, T. R. Brummelkamp, J. A. Wojcechowskyj, A haploid genetic screen identifies heparan sulfate proteoglycans supporting Rift Valley fever virus infection. J. Virol. 90, 1414–1423 (2016).

62. K. H. D. Crawford, R. Eguia, A. S. Dingens, A. N. Loes, K. D. Malone, C. R. Wolf, H. Y. Chu, M. A. Tortorici, D. Veesler, M. Murphy, D. Pettie, N. P. King, A. B. Balazs, J. D. Bloom, Protocol and Reagents for Pseudotyping Lentiviral Particles with SARS-CoV-2 Spike Protein for Neutralization Assays. Viruses. 12 (2020), doi:10.3390/v12050513.

63. A. C. Walls, M. C. Miranda, M. N. Pham, A. Schäfer, A. Greaney, P. S. Arunachalam, M.-J. Navarro, M. A. Tortorici, K. Rogers, M. A. O’Connor, L. Shireff, D. E. Ferrell, N. Brunette, E. Kepl, J. Bowen, S. K. Zepeda, T. Starr, C.-L. Hsieh, B. Fiala, S. Wrenn, D. Pettie, C. Sydeman, M. Johnson, A. Blackstone, R. Ravichandran, C. Ogohara, L. Carter, S. W. Tilles, R. Rappuoli, D. T. O’Hagan, R. Van Der Most, W. C. Van Voorhis, J. S. McLellan, H. Kleanthous, T. P. Sheahan, D. H. Fuller, F. Villinger, J. Bloom, B. Pulendran, R. Baric, N. King, D. Veesler, Elicitation of broadly protective sarbecovirus immunity by receptor-binding domain nanoparticle vaccines. bioRxivorg (2021), doi:10.1101/2021.03.15.435528.

64. J. Gregson, S. Y. Rhee, R. Datir, D. Pillay, C. F. Perno, A. Derache, R. S. Shafer, R. K. Gupta, Human immunodeficiency virus-1 viral load is elevated in individuals with reverse-transcriptase mutation M184V/I during virological failure of first-line antiretroviral therapy and is associated with compensatory mutation L74I. J. Infect. Dis. 222, 1108–1116 (2020).

65. X. Ou, Y. Liu, X. Lei, P. Li, D. Mi, L. Ren, L. Guo, R. Guo, T. Chen, J. Hu, Z. Xiang, Z. Mu, X. Chen, J. Chen, K. Hu, Q. Jin, J. Wang, Z. Qian, Characterization of spike glycoprotein of SARS-CoV-2 on virus entry and its immune cross-reactivity with SARS-CoV. Nat. Commun. 11, 1620 (2020).

